# Determining Reliable Neural Targets for enhancing Motor Sequence Learning: A preparatory study for focal tDCS

**DOI:** 10.64898/2026.06.23.733856

**Authors:** Asad Khan, Michael Burke, Erhan Genc, Harleen Chhabra, Min-Fang Kuo, Michael A. Nitsche

## Abstract

**Background:** Motor sequence learning (MSL) underlies the acquisition of everyday skilled movements, and transcranial direct current stimulation (tDCS) has shown promise for enhancing it, although reported effects vary considerably. Before such interventions can be applied and interpreted reliably, the behavioural and neural signatures of the underlying task should be shown to be robust. This study therefore characterised an fMRI-based Serial Reaction Time Task (SRTT) protocol and evaluated the consistency of its behavioural and fMRI-based neural signatures in repeated measurements to identify potential network targets for tDCS.

**Methods:** Twenty healthy young-to middle-aged adults (10 females; mean age 24, SD 4) performed the SRTT during 3T fMRI in two sessions about one week apart, each using one of two matched, non-overlapping task versions in counterbalanced order. We examined block-wise behaviour, task-related and sequence-specific brain activation (general task activation relative to baseline and the SEQ > RND contrast), and functional connectivity using two complementary approaches: the task and temporal dynamics of seed-based connectivity (SBC) within an a priori-defined motor network (M1, SMA, PMC, putamen, and cerebellum), and whole-brain generalised psychophysiological interaction (gPPI) for the SEQ > RND contrast.

**Results:** Reaction time decreased steadily across sequence blocks and returned toward baseline during random blocks, confirming implicit sequence learning; this pattern was consistent across both sessions and both task versions. Test–retest reliability was excellent for random-block reaction time (ICC ≈ .80 – .85), and modest for sequence blocks (ICC ≈ .29 – .53). General task activation (sequence and random performance relative to baseline) engaged the expected sensorimotor network, including the primary motor cortex (M1), premotor cortex, and SMA, with M1 more strongly engaged during sequence than random blocks. The sequence-specific contrast (SEQ > RND) additionally revealed six clusters spanning the left superior parietal cortex, bilateral supplementary and cingulate motor areas, bilateral secondary somatosensory cortex, and left ventral visual cortex together with cerebellar Crus I. The a priori defined motor network was engaged across all task phases and was anchored throughout by strong M1–SMA coupling, with overall network coupling following a non-monotonic course across the session, consistent with successive stages of resource allocation, consolidation, and automatisation. The gPPI analysis identified four sequence-specific connectivity clusters, dominated by a right-lateralised visuomotor network and a somatosensory–visual integration channel.

**Conclusions:** Together, these findings indicate a two-component architecture for SRTT performance: a left hemispheric cortico–striato–cerebellar network that supports performance and acquisition of the sequence, and a right hemispheric network associated with monitoring its predictable structure. Within this protocol, the SRTT produced robust and stable group-level signatures of implicit motor learning, yielding well-defined network targets for future tDCS interventions..

## 1 Introduction

Motor learning, the process by which practice produces lasting improvements in skilled movement, is fundamental to everyday function across the lifespan. One of its most studied forms is motor sequence learning (MSL), the acquisition of an ordered series of movements that, with practice, are executed faster, more accurately, and with progressively less attentional demand. Like other forms of motor learning, MSL declines with age (Leuk et al., 2025), contributing to reduced independence in older adults. Implicit MSL is commonly assessed using the Serial Reaction Time Task (SRTT; Nissen & Bullemer, 1987), in which participants respond to visual cues that follow either a hidden repeating sequence or a random order of stimuli. Learning is inferred from progressively faster and more accurate responses to the sequence, together with a return toward baseline speed when random material is reintroduced (Robertson, 2007; Schwarb & Schumacher, 2012).

The physiological mechanisms underlying MSL have been studied extensively in humans using neuroimaging. SRTT learning engages a distributed cortico–striato–cerebellar network spanning the primary motor cortex (M1), the premotor and supplementary motor areas (SMA), the parietal cortex, the basal ganglia, the thalamus, and the cerebellum (Doyon & Benali, 2005; Hardwick et al., 2013; Hikosaka et al., 2002; Janacsek et al., 2020). These regions are thought to contribute differentially across learning stages: M1 and subcortical structures are most engaged during early learning, whereas premotor and parietal cortices, together with the SMA, become more important in later stages and in representing the abstract sequence independently of the effectors used to perform it (Grafton et al., 2002; Honda et al., 1998; Muellbacher et al., 2002; Nachev et al., 2008).

More recent imaging work has refined, and in part challenged, this picture. Some studies have questioned whether M1 carries a genuinely sequence-specific representation, attributing much of its activation to movement execution rather than to learning of the sequence itself (Berlot et al., 2020). Other recent studies, however, stress a critical role for M1, particularly in early sequence learning, emphasising the progressive reconfiguration of its cortico-cortical and cortico-subcortical coupling as the sequence is acquired (Chen et al., 2026; Hamano et al., 2020; Wiestler & Diedrichsen, 2013). Convergent evidence comes from non-invasive brain stimulation: anodal tDCS applied over M1 during learning facilitates implicit MSL and its consolidation (Nitsche et al., 2003; Reis et al., 2009), whereas stimulation of premotor and prefrontal control sites during early learning was without effect, supporting the specificity of M1 contribution at this stage (Nitsche et al., 2003).

Because fMRI activation and connectivity are correlational measures, they can establish which regions are engaged during task performance but cannot on their own determine the causal contribution of a given area to learning. Establishing causality requires intervening directly in the system, which can be achieved with non-invasive brain stimulation such as tDCS. tDCS is well suited for this purpose because it engages physiological mechanisms thought to underlie motor memory formation: anodal tDCS over M1 induces long-term-potentiation (LTP)-like plasticity through increased glutamatergic NMDA-receptor activity and reduced GABAergic inhibition (Nitsche & Paulus, 2000; Stagg et al., 2018), synaptic processes involved in the formation and consolidation of motor skills (Rioult-Pedotti et al., 2000). Targeting a candidate area during task performance therefore allows to determine its causal and physiological role in MSL.

Reported tDCS effects on MSL are, however, variable. A substantial part of this variability arises from differences in stimulation protocols across studies with respect to target determination, timing, montage, and dosing, including dosage-dependent, non-linear physiological effects and is most pronounced for conventional, non-focal montages with non-individualised electrode placement (Buch et al., 2017; Hashemirad et al., 2016; Woods et al., 2016). At the same time, the behavioural and neural signatures of the SRTT differ across studies, and both behavioural and task-based fMRI measures often show low test–retest reliability (Elliott et al., 2020; Judd et al., 2024; Smith et al., 2005). This is decisive for crossover and longitudinal interventions, which depend on the stability of the targeted measures across repeated testing (Meinzer et al., 2024). For a planned tDCS study, the group-level behavioural and imaging signatures must therefore be shown to be reproducible across sessions and task versions, and their reliability quantified, before active stimulation is applied.

Characterising the physiological foundation of the task in this way requires knowing which regions are activated during performance, how the relevant network is connected, and how that connectivity is specific to sequence processing. To this end, we combined three complementary fMRI analyses: whole-brain BOLD activation, which localises the regions recruited by general task performance and by sequence processing specifically; seed-based connectivity (SBC) of an a priori defined motor network, which describes how that network is configured within each task condition; and generalised psychophysiological interaction (gPPI), which identifies, in an exploratory whole-brain manner, the connections specifically modulated by sequence learning (Di et al., 2021; McLaren et al., 2012). Together, these analyses define the functional network engaged by the task and identify candidate targets for subsequent focal stimulation.

Specifically, the present study was designed to establish an SRTT-based MSL protocol that yields clear and stable group-level behavioural and neural signatures, can be performed inside the MRI scanner, and provides a validated basis for a subsequent focal-tDCS study. Healthy young-to middle-aged adults performed the SRTT during fMRI in two sessions using two matched, non-overlapping task versions. The study had the following aims: (i) to verify the group-level behavioural signature of implicit MSL (ii) to map both the general task activation and the sequence-specific activation during task (iii) to characterise the functional connectivity architecture of the task, and (iv) to assess the group effects and test–retest reliability of these behavioural, activation and connectivity measures. Together, these measures aimed to establish the suitability of the paradigm for investigating tDCS effects in a subsequent sham-controlled, intra-scanner crossover study.

## 2 Materials and Methods

This study reports findings from the preparatory phase of a multicentre, double-blind, sham-controlled crossover trial examining the behavioural and neural effects of focal tDCS across motor and cognitive domains (https://www.memoslap.de/de/forschung/). The aim of this phase was to validate task paradigms targeting four functional domains, visual-spatial, language, executive, and motor, and to assess their reliability, in order to identify candidate stimulation targets for subsequent phases of the trial. The study was pre-registered on the Open Science Framework (OSF), and the protocol is publicly available via OSF Registries (osf.io/t37u2). An overview of the intra-scanner task design and the experimental procedures of this study is provided in Figure 1.

**Figure 1.**
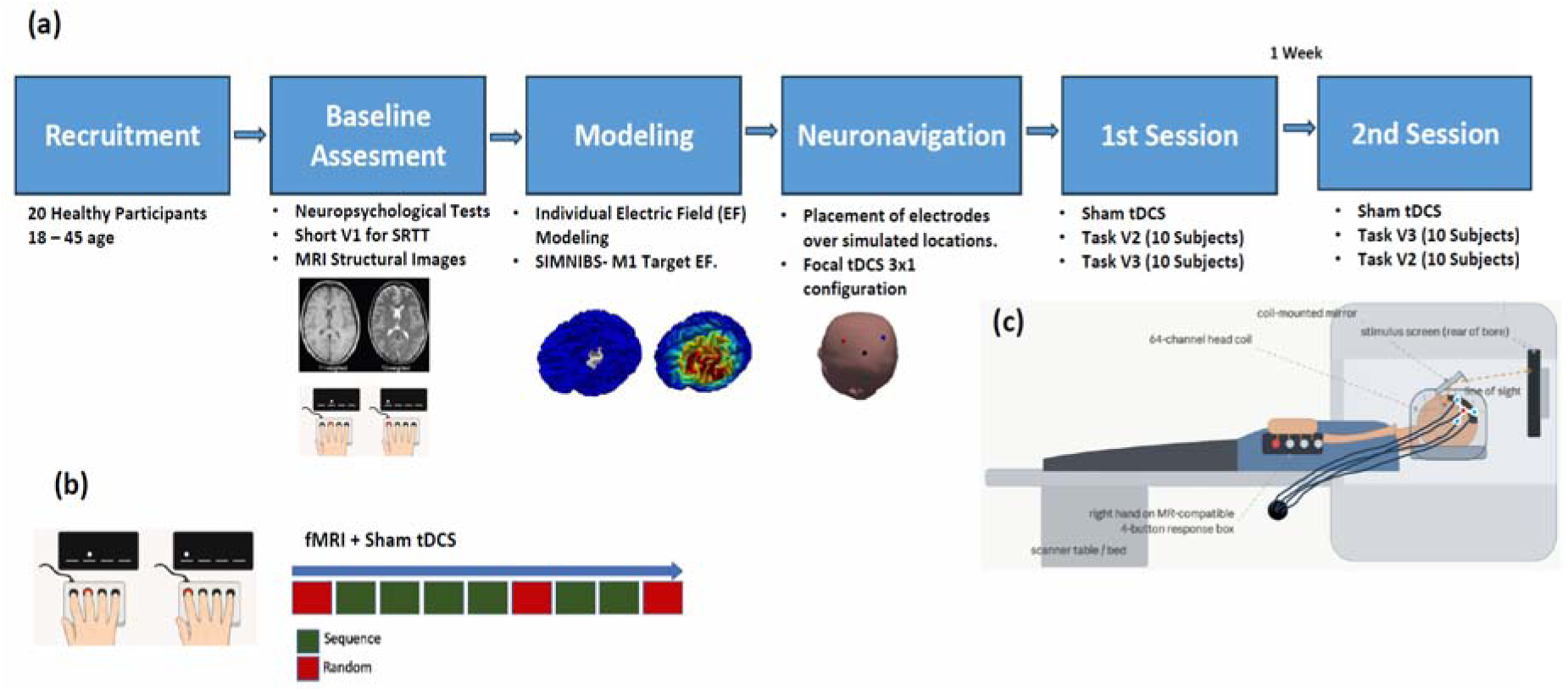
(A) Flowchart of the experimental procedure. Twenty healthy participants (aged 18–45) underwent baseline assessments, including neuropsychological testing, a short version of the SRTT task, and structural MRI (T1 and T2). Individualised electric field modelling was performed using SimNIBS, targeting the left M1 followed by neuronavigation-guided focal transcranial direct current stimulation (tDCS) (3 × 1 montage). Participants completed two task-based fMRI sessions, separated by 1 week. All participants performed both Task V2 and Task V3, with task order counterbalanced across sessions (i.e., 10 participants completed Task V2 in Session 3 and Task V3 in Session 4, whereas the other 10 participants completed the tasks in the reverse order). Each session included brief sham tDCS and task performance inside the scanner. (B) Block design of the fMRI task. Each task session consisted of six sequence (learning) blocks (2-5, 7 and 8, marked in Green) (SRTT task) and three Random (Control) blocks (1, 6 and 9, marked in Red). (C) Responses were made using an MRI-compatible response box. fMRI was measured while the subject was shown the SRTT task on a coil mounted mirror inside the MRI in supine position.

### 2.1 Experimental Design

#### 2.1.1 Participants

Twenty healthy young-to middle-aged adults (range 18–45) participated (10 females; mean age = 24; standard deviation [SD] = 4). All participants were non-smokers, right-handed, as evaluated by the Edinburgh Handedness Inventory (Oldfield, 1971); they were native German speakers and reported no history of neurological or psychiatric conditions. Participants also had no history of alcoholism, drug use, or contraindications for MRI or tDCS (Antal et al., 2017). A physician examined all volunteers for exclusion criteria for non-invasive electrical brain stimulation. Volunteers participating in the study signed an informed consent form and received financial compensation.

#### 2.1.2 Baseline Assessments

Participants underwent an initial telephone screening to confirm their eligibility, including screening for MRI- and tES-specific inclusion/exclusion criteria (Antal et al, 2017). Following this, a comprehensive battery of questionnaires, neuropsychological tests, and motor-dexterity assessments was administered to characterise the participants’ baseline status. Caffeine intake and other factors potentially interfering with cortical excitability (e.g., strenuous physical activity within 48 h prior to a session) were controlled for before each experimental session.

Questionnaires: Demographic information (age, sex, educational level) was collected with a demographic questionnaire. Depressive symptoms were screened with the German version of Beck’s Depression Inventory (BDI-II; Kühner et al., 2007), and chronotype was assessed with the German version of the Morningness–Eveningness Questionnaire (D-MEQ; Horne & Östberg, 1976). The amount of habitual physical activity was obtained with the Lüdenscheider Activity Questionnaire (Höltke & Jakob, 2002), reported as Total and Sports subscale scores.

Neuropsychological assessment: Verbal episodic memory was assessed with the German Auditory Verbal Learning Test (Verbal Learning and Memory Test, VLMT; Müller et al., 1997), reported as total learning (sums of trials DG1–DG5), delayed recall (DG7), and recognition (DG8). Verbal working memory was assessed with the Digit Span backward and spatial working memory with the Corsi Block Tapping Test, forward and backward (Wechsler, 1987). Executive functioning was assessed with the Stroop Colour–Word Test (word reading, colour naming, and interference conditions; Van der Elst et al., 2006). Perceptual speed was assessed with the Digit Symbol Substitution Test (DSST; Wechsler, 1981), and verbal intelligence / vocabulary knowledge with the German multiple-choice vocabulary test (MWT-B; Lehrl, 2018).

Motor-dexterity assessment: Fine and gross motor function was characterised with two standardised instruments. The SCAFI (Schmitz-Hübsch et al., 2008) provided measures of gait speed (8 m walking time) and manual dexterity (nine-hole peg-board completion times for the right and left hands). The Motor Performance Series (Motorische Leistungsserie, MLS; Schuhfried GmbH, Mödling, Austria) provided fine-motor measures of aiming, steadiness, line tracking, and tapping, together with the standardised MLS factor scores for manual unrest/tremor (Factor 2), precision (Factor 3), arm–hand speed (Factor 5), and wrist–finger speed (Factor 6).

Study-specific assessment: Participants also completed a short version (Version V1) of the SRTT (three blocks), both as practice for the main intra-scanner task and to classify them as “high” or “low” performers in the learning task. This classification is potentially relevant for tDCS outcomes (Meinzer et al., 2013; Perceval et al., 2020) and and served as a predictor of effects in subsequent active stimulation phases of the larger study. Participants’ characteristics are provided in Table 1.

**TABLE 1.**
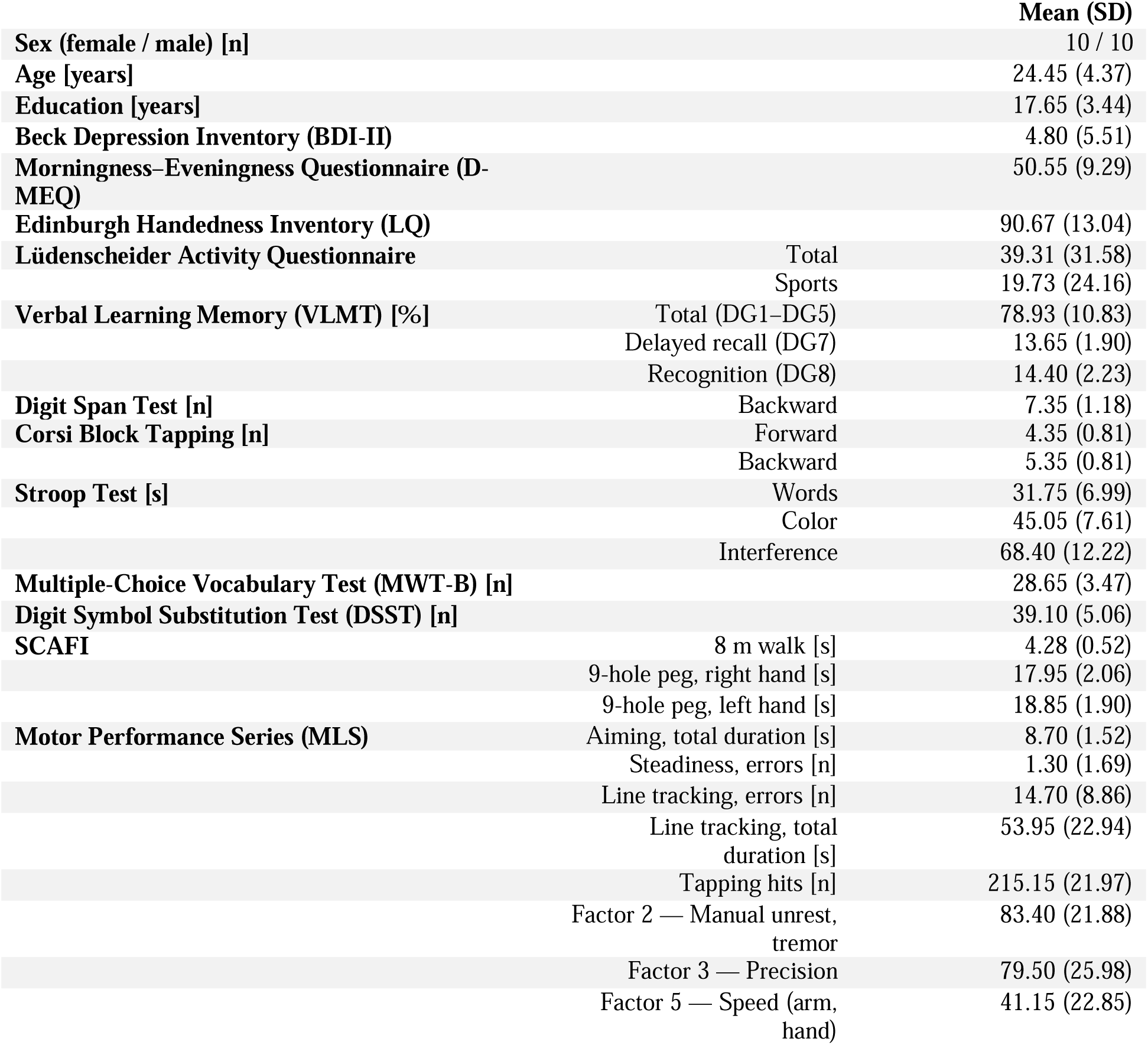
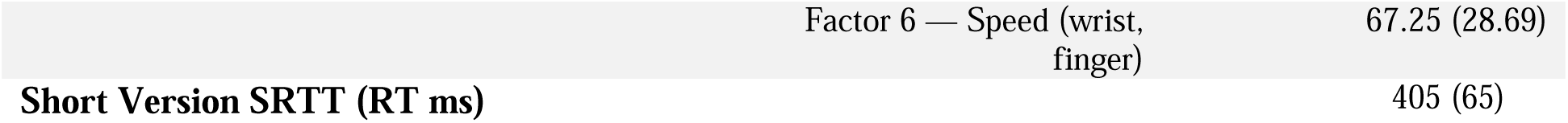
Descriptive statistics of demographic, neuropsychological, and motor-performance data of the study sample.

#### 2.1.3 SRTT Task Design

The Serial Reaction Time Task (SRTT) is a motor sequence learning task. Performance on this task is associated with increased activity and cortical excitability of the motor, premotor, and supplementary motor areas, and early learning prominently involves the primary motor cortex (Hardwick et al., 2013; Hikosaka et al., 2002; Janacsek et al., 2020; Honda et al., 1998; Muellbacher et al., 2002; Hamano et al., 2020; Nitsche et al., 2003).

In this task, sequence motor learning is primarily indicated by a reduction of reaction time when participants press the appropriate button on a response box after the presentation of a visually cued stimulus on a computer screen (Figure 1(b)).

A custom-made response box with four response keys anatomically aligned to the right hand was used to record performance. The four fingers involved in performance were the index finger for the first button, the middle finger for the second button, the ring finger for the third button, and the little finger for the fourth button (buttons from left to right). Four horizontal lines representing the keys were presented on a computer screen. Participants were instructed to press the corresponding key with the correct finger as fast and accurately as possible when a stimulus (white dot) appeared above the corresponding line. The next trial appeared 500 ms following a button press, independent of a correct or incorrect response. The task consisted of 9 blocks, each with 120 trials. Blocks 1, 6 and 9 were random blocks, in which the trials were presented in a pseudo-random order. Blocks 2–5 and 7–8 displayed a 12-trial sequence of stimuli repeated 10 times in each block, which prompted implicit motor sequence learning (Robertson & E. M., 2007, Nitsche et al., 2003, Kuo et al., 2008; Nitsche et al., 2010). Blocks were separated by self-paced breaks and participants were not informed about the sequences. In order to control for use-dependent sequence learning over sessions, two different sequences of the task with no overlapping parts and comparable difficulties were presented in the two sessions in counterbalanced order; these are referred to throughout the manuscript as Task Version 2 (V2) and Task Version 3 (V3). Participants were not informed about the repeating sequence. At the end of the session, they were asked whether they had noticed a sequence and, if so, to write it down, in order to assess explicit learning. Participants who reported having noticed the sequence, and were able to reproduce it, were classified as having developed explicit knowledge of the task, and their data were excluded from the final analysis.

#### 2.1.4 Focal Sham tDCS

This study aimed to characterise the brain regions and networks engaged during implicit motor sequence learning in the Serial Reaction Time Task (SRTT), to identify candidate targets for focal tDCS interventions. The main objective of the present study was to establish a stable group-level signature of behavioural performance, and fMRI-based activation and connectivity. Sham tDCS was applied in both sessions to develop and evaluate the practical workflow for future combined tDCS-fMRI studies. This workflow included neuronavigated electrode placement, delivery of stimulation inside the MRI scanner, task performance during image acquisition, and optimisation of the fMRI acquisition protocol. The sham condition also provided a suitable control condition for later phases of the broader project, in which active stimulation was tested (Meinzer et al., 2024).

Stimulation was delivered using a focal 3 × 1 tDCS (Figure 1 (c)) configuration connected to a multi-channel direct current stimulator. Sham tDCS was applied during the functional MRI sessions with MRI-compatible components, including a multi-channel Starstim stimulator (NeuroElectrics, Barcelona, Spain), circular conductive rubber electrodes measuring 2 cm in diameter, and MRI-compatible cables. (neuroConn GmbH, Ilmenau, Germany). Further technical details are provided by Niemann et al. (2024). The sham protocol consisted of a 10-s ramp-up phase, 30 s of stimulation at 2 mA, and a 10-s ramp-down phase. This approach was intended to imitate the initial sensation of active tDCS without producing sustained changes in cortical excitability (Nitsche et al., 2008). To ensure comparability with the later experimental phases, sham stimulation was applied with the same montage and current intensity planned for the active tDCS arms of the overarching project, and tDCS was applied immediately before task performance to avoid interference with performance.

The left primary motor cortex (M1) was chosen as the target region on the basis of converging neuroimaging and stimulation evidence identifying the contralateral primary motor cortex as a central node of implicit motor sequence learning for the right hand (Doyon & Benali, 2005; Hardwick et al., 2013; Hikosaka et al., 2002; Janacsek et al., 2020). Nitsche et al., 2003; Reis et al., 2009; Stagg et al., 2011).

The anode was positioned over the individually identified left M1 hand-knob region and surrounded by three evenly spaced cathodes, with consistent spacing across participants. To improve blinding, a topical anaesthetic cream (EMLA®, 2.5% lidocaine, 2.5% prilocaine) was applied to all electrode sites (Garofalo et al., 2021).

Cathode spacing was determined based on pilot electric-field simulations to ensure consistent received stimulation intensity at the target of 0.2 mA across participants and MeMoSLAP subprojects (Abdelmotaleb et al. 2025). Electrode positioning was standardised using a 3D-printed thermoplastic spacer with a 3 cm radius, which defined the fixed distance between the central anode and the surrounding cathodes (Niemann et al., 2024).

For montage individualisation, the optimisation procedure was designed to robustly focalise the electric field over the participant-specific target region while minimising current spread to adjacent cortical areas. Subject-specific tetrahedral head models were generated from T1- and T2-weighted anatomical images using the charm pipeline, followed by tissue segmentation and electric-field modelling in SimNIBS v4 (simnibs.org; Thielscher et al., 2011; Puonti et al., 2020).

While only sham tDCS was administered in the present study to characterise behavioural and neural signatures of SRTT performance, identical modelling and electrode-positioning procedures were implemented to mirror the individualised stimulation protocol planned for the subsequent active tDCS phases of the broader MeMoSLAP study.

#### 2.1.5 MRI Acquisition

All MRI data were acquired on a 3.0 T Siemens MAGNETOM PRISMA system (Siemens Healthineers, Erlangen, Germany) at the Leibniz Research Centre for Working Environment and Human Factors, Dortmund, Germany, fitted with a 64-channel head–neck coil. Functional runs used a multiband echo-planar imaging (EPI) protocol provided by the Center for Magnetic Resonance Research (CMRR, University of Minnesota). This protocol relies on simultaneous multislice (SMS) acceleration and is tuned for blood oxygenation level-dependent (BOLD) sensitivity; by shortening the per-slice repetition interval, it permits a high sampling rate while preserving whole-brain coverage (Setsompop et al. 2012; Xu et al. 2013).

For the task runs, volumes were sampled with a 110 × 110 acquisition matrix, an in-plane resolution of 2 × 2 mm^2^, and a 2 mm slice thickness with no interslice gap. The governing parameters were a repetition time (TR) of 1000 ms, echo time (TE) of 31 ms, flip angle of 60°, field of view (FOV) of 220 mm, and a multiband acceleration factor of 6. Each run lasted approximately 20 minutes and produced 1200 functional volumes, with phase encoding applied along the anterior-to-posterior (AP) axis.

To limit geometric distortion and signal drop-out, echo spacing was set to 540 µs, and slices were collected in interleaved order to attenuate motion-related effects. Field maps for distortion correction were obtained separately by reversing the phase-encoding polarity. High-resolution anatomical scans were acquired at 0.9 mm isotropic resolution: a T1-weighted volume (TR = 2700 ms, TE = 3.7 ms, TI = 1090 ms, flip angle = 9°), with selective water excitation used for fat suppression, and a T2-weighted volume at the same resolution (TR = 2500 ms, TE = 349 ms).

### 2.2 Statistical Analysis

#### 2.2.1 Behavioural Data Analysis

Behavioural performance was analysed at the block level for four dependent variables: means of absolute reaction time (RT, ms); means of normalised RT (each block’s RT expressed as a ratio relative to Block 1); RT variability (within-block standard deviation of RT); and accuracy (proportion of correct responses). Two separate complementary two-way repeated-measures ANOVAs were conducted rather than a single three-way model, because these addressed two distinct questions. The first (Session analysis) compared the two sessions (factor Session, session1, session 2) in order to exclude session-specific performance differences, which would reflect effects of repeated exposure to the task rather than an effect of stimulation. The second (Task Version analysis) compared the two task versions (Task V2 vs. Task V3), which were counterbalanced across participants, to confirm that the two versions yielded comparable performance. Each of the two ANOVAs (Sessions and Task Versions) was conducted separately for every dependent variable (absolute RT, normalised RT, RT variability, and accuracy). The Session analysis was a 2 × 9 repeated-measures ANOVA with the within-subject factors Session (two levels: Session 1, Session 2) and Block (nine levels: Blocks 1–9). Likewise, the Task Version analysis was also a 2 × 9 repeated-measures ANOVA with the within-subject factors Block (nine levels: Blocks 1–9) and Task Version (two levels: V2, V3). For the ANOVA’s Mauchly’s test sphericity was conducted, and the Greenhouse Geisser correction was applied when necessary and sizes are reported as partial eta squared (ηp^2^).

Post-hoc tests were conducted separately within each ANOVA in case of significant ANOVA effects. Within each analysis, paired t-tests with Fisher’s least significant difference compared each block against Block 1. To test directly if blocks with random stimuli differed from sequence blocks with respect to RT, and thus explore specific sequence effects on performance, another ANOVA was conducted for the RT differences between Block 5-6 and Block 6-7. Additional paired t-tests were conducted on the change scores ΔB5–B6 and ΔB6–B7 in case of significant ANOVA results. Effect sizes for these post-hoc comparisons are reported as Cohen’s d. Block-wise test–retest reliability was assessed between Sessions 1 and 2 for reaction time and accuracy. Two forms of the intraclass correlation coefficient were computed for each block (1–9): ICC(2,1), which indexes absolute agreement and penalises session-to-session mean shifts; and ICC(3,1), which indexes consistency and assesses whether participants retain a stable rank ordering across sessions independently of global performance shifts (Koo & Li, 2016; McGraw & Wong, 1996; Shrout & Fleiss, 1979). Comparing these two parameters allowed systematic session-related shifts (such as practice effects) to be distinguished from true instability. Ninety-five percent confidence intervals for each ICC were estimated by bootstrap resampling with 1000 iterations. Reliability was interpreted against conventional thresholds of ICC ≥ .75 (good) and ICC ≥ .50 (moderate) (Koo & Li, 2016).

#### 2.2.2 fMRI Data Analysis

Preprocessing was carried out with fMRIPrep (Esteban et al., 2019), a Nipype-based preprocessing pipeline. For each participant and session, functional images were corrected for slice timing, realigned to estimate and correct head motion, distortion correction, co-registered to the participant’s anatomical image, and normalised to MNI152 standard space. The preprocessed and normalised functional images were then imported into SPM12 (Wellcome Trust Centre for Neuroimaging, London, UK). Spatial smoothing was applied in SPM12 using a 6 mm full-width at half-maximum Gaussian kernel before statistical analysis.

Functional imaging data were analysed using the general linear model (GLM) with a block-design approach. A separate first-level model was specified for each participant and session. Each model included nine experimental blocks, consisting of six sequence blocks and three random blocks. Blocks were modelled as boxcar functions using the duration of each block and were convolved with the canonical haemodynamic response function. The six rigid-body motion parameters estimated by fMRIPrep were included as nuisance regressors to account for motion-related variance. A rest condition was also modelled as a separate regressor after the last block which acted as the baseline from which activation map contrasts were created compared to the task conditions. Throughout the paper this rest condition will be called baseline.

Three whole-brain group-level contrasts for BOLD activity were computed, (i) SEQ > Baseline, (ii) RND > Baseline, and (iii) the direct sequence-specific contrast SEQ > RND.

Based on the results of the test–retest analyses (see Section 2.2.4), these contrasts were computed on whole-group data.

The SEQ > RND contrast was used as the main test of sequence-specific neural activity, because it removes activation related to general task execution that is common to both sequence and random blocks. Statistical maps were thresholded using a voxel-wise cluster-forming uncorrected threshold of p < .001. Clusters were considered significant at p < .05 after cluster-level family-wise error correction. Activations were displayed on the MNI152 template as axial slices and 3D cortical surface renderings.

Peak regions were labelled using the Harvard–Oxford cortical and subcortical structural atlas. The Automated Anatomical Labelling atlas was used when peaks did not reach the 25% probability threshold in the Harvard–Oxford atlas or when peaks were located in cerebellar regions not covered by in that atlas. These labels are marked with an asterisk in the activation Tables.

#### 2.2.3 Connectivity analysis

##### Seed-based connectivity (SBC)

Seed-based functional connectivity (SBC) and the gPPI analysis described below are both seed-based approaches: SBC quantifies condition-wise coupling of the predefined seeds, whereas gPPI tests show how coupling is modulated by the task contrast. The SBC connectivity was analysed within a predefined motor network consisting of five regions of interest: left primary motor cortex (L_M1), left supplementary motor area (L_SMA), left premotor cortex (L_PMC), left putamen (L_Putamen), and right cerebellum (R_Cerebellum). These regions were selected because they are part of the motor network involved in right-hand MSL performance. (Tzvi et al., 2014, 2015; Liebrand et al., 2020). The left cortical and subcortical regions are contralateral to the responding hand, while the right cerebellum was included because of its functional connection to right-hand motor control through crossed cerebellar pathways (Bernard & Seidler, 2013; Liebrand et al., 2020; Stoodley & Schmahmann, 2009).

For each participant, session, and task condition, the average BOLD signal was extracted from each of the pre-defined ROIs. Functional connectivity between ROI pairs was then calculated using Fisher-z-transformed partial correlations. Before estimating connectivity, nuisance signals from white matter, cerebrospinal fluid, and six motion parameters were regressed out, and the data were band-pass filtered (0.008 -0.09 Hz). These analyses were performed using the standard seed-based connectivity pipeline implemented in the CONN toolbox (Whitfield-Gabrieli & Nieto-Castañón, 2012).

SBC was was estimated separately for five task conditions: Initial RND, corresponding to Block 1 before sequence exposure; Early SEQ, including Blocks 2–3; Middle SEQ, including Blocks 4–5; Late SEQ, including Blocks 7–8; and Interleaved RND, including Blocks 6 and 9. Data from Sessions 1 and 2 were combined for this analysis. As detailed in Section 2.2.4, this pooling was justified by the absence of group-level differences between sessions and task versions, established by paired-samples t-tests (Supplementary Material).

By including Block 1 as an initial pre-sequence random baseline, the present design allows direct comparison between random blocks before and after sequence learning. At the group level, connectivity results for each task condition were analysed across Sessions 1 and 2 using an omnibus F-test. Statistical inference was performed in CONN using its network-based statistics approach. A connection-level threshold of p < .05, uncorrected and two-sided, was first applied. Significant network connections were then identified using a cluster-level threshold of p < .05 with FDR correction, based on the omnibus multivariate pattern analysis test. For each significant cluster, individual connections were also examined using family-wise error correction at the connection level (p-FWE < .05). Because the analysis included only five ROIs, resulting in 10 possible pairwise connections, the FDR- and FWE-corrected results were very similar. All positive connections reported in the results survived both correction approaches.

##### Generalised psychophysiological interaction (gPPI)

Task-related changes in functional connectivity were analysed using generalised psychophysiological interaction analysis, gPPI (McLaren et al., 2012). The analysis focused on the SEQ > RND contrast and combined data from Sessions 1 and 2. A total of 132 cortical and subcortical regions from the Harvard–Oxford atlas was included, resulting in in 17,292 ROI-to-ROI connections.

Functional network clusters were defined using hierarchical clustering based on the ROI-to-ROI connectivity patterns. These clusters were then tested using an omnibus F-test based on multivariate pattern analysis, with cluster-level FDR correction. For clusters that survived this correction, individual connections were further examined using family-wise error correction at the connection level (p-FWE < .05). Positive T-values indicate stronger connectivity during sequence blocks compared with random blocks, whereas negative T-values indicate weaker connectivity during sequence blocks.

#### 2.2.4 Reproducibility and reliability analysis

Two complementary properties of each measure were assessed: group-level reproducibility across the two sessions and the two counterbalanced task versions, and individual-level test–retest reliability. These address different questions and do not necessarily coincide. Group-level reproducibility addresses the question whether the mean effect is stable across repetitions and task versions, and the absence of such differences justifies pooling the data. Individual-level reliability, quantified by the intraclass correlation coefficient (ICC), asks whether participants retain a stable rank ordering across measurements. Because the SRTT evokes a highly consistent, near-universal learning effect, between-subject variance is small, and individual-level reliability can be only poor to moderate even when the group-level effect is robust and stable, a pattern known as the reliability paradox (Hedge et al., 2018). Difference contrasts such as SEQ > RND are expected to be the least reliable because SEQ and RND are highly correlated across participants, thus contrasting these removes most of the stable between-participant variance while the independent measurement errors of the two conditions add, lowering the intraclass correlation of the difference (Hedge et al., 2018). We therefore quantified both types of analyses for all three data modalities (behaviour, BOLD activation, and connectivity) and based the decision to pool sessions and versions on the group-level results rather than on ICC.

##### Behaviour

Group-level reproducibility was tested with the 2 × 9 repeated-measures ANOVAs and paired t-tests described in Section 2.2.1 (Session and Task Version analyses). Individual-level reliability was quantified with ICC(2,1) (absolute agreement) and ICC(3,1) (consistency) for each block, as described in that section.

##### BOLD activation

Group-level reproducibility was tested with whole-brain paired-samples t-tests on the first-level contrast images (SEQ > Baseline, RND > Baseline, and SEQ > RND), comparing Session 1 with Session 2 and Task V2 with Task V3 (voxel-wise p < .001, cluster-level FWE p < .05). Individual-level reliability was quantified at three spatial scales using independent, anatomically defined regions rather than clusters derived from the contrasts, to avoid circularity: a whole-image intraclass correlation (I2C2; Shou et al., 2013), voxel-wise ICC, and atlas-based ROI ICC, each computed separately for the SEQ, RND, and SEQ > RND maps. ICC(2,1) was reported as the primary index alongside ICC(3,1). Region-level equivalence between sessions and versions was additionally evaluated with two one-sided tests (TOST) and JZS Bayes factors, to provide positive evidence for the absence of differences.

##### Connectivity

For the seed-based connectivity (SBC) network, group-level reproducibility was tested with per-edge paired-samples t-tests comparing sessions and versions within each task phase, with false-discovery-rate correction across the ten edges, complemented by TOST and JZS Bayes-factor equivalence tests; individual-level reliability was quantified per edge with ICC(3,1) and ICC(2,1). For the exploratory gPPI analysis, the 17,292 directed edges make per-edge testing statistically inappropriate; reproducibility of the whole SEQ > RND connectivity pattern was therefore assessed with the image intraclass correlation (I2C2) and the spatial correlation between the session- and version-specific group matrices, evaluated against a permutation null, and the full distribution of per-edge ICCs was summarised. Detailed reproducibility and reliability results for activation and connectivity are reported in the Supplementary Material.

## 3 Results

### 3.1 Reproducibility and reliability of SRTT measures across sessions and task versions

We first established that the SRTT measures were reproducible across the two sessions and the two counterbalanced task versions, which justifies the pooling used in all subsequent analyses, and we characterised their individual-level reliability. Consistent with the known behaviour of this paradigm, the data showed a clear dissociation between stable group-level effects and lower individual-level reliability.

#### Group-level reproducibility

At the group level, sessions and task versions did not differ. Behaviourally, neither session, task version, nor their interaction reached significance for any measure (Section 3.2). For BOLD activation, whole-brain paired-samples t-tests of the SEQ > Baseline, RND > Baseline, and SEQ > RND contrasts revealed no clusters surviving correction between Session 1 and Session 2; the only difference between task versions was a single small cluster in left inferior frontal / ventral premotor cortex for the SEQ contrast, which, given the counterbalancing of V2 and V3 across sessions, does not bias the pooled analyses. For connectivity, the seed-based network showed no systematic session or version differences (2 of 50 edge-tests differed between sessions and none between versions, after correction), and the exploratory gPPI SEQ > RND pattern did not differ between sessions or versions. These results justify pooling the two sessions and the two versions on the basis of group-level equivalence (Supplementary Material).

#### Individual-level reliability

At the individual level, test–retest reliability was lower, as expected. Behaviourally, reaction time in the random blocks was highly reliable (ICC ≈ .80–.85), whereas sequence-block reaction time was only modestly reliable (ICC ≈ .29–.53; Section 3.3). For BOLD activation, the SEQ and RND maps showed moderate reliability (whole-image I2C2 ≈ 0.47 and 0.46; median atlas-ROI ICC ≈ 0.53 and 0.46), whereas the SEQ > RND difference map was close to zero (I2C2 ≈ 0.08). For connectivity, edge-level reliability of the seed-based network was low overall and increased with the number of pooled blocks (median ICC rising from near zero for individual phases to ≈ 0.36 across all blocks), while the gPPI SEQ > RND difference pattern showed essentially no individual-level reliability (I2C2 ≈ 0.01).

Thus the SRTT and its imaging correlates produced reproducible, stable group-level effects together with low-to-moderate individual-level reliability, and the lowest reliability occurred precisely for the difference contrasts (SEQ > RND) in which the shared signal is removed. This pattern is the expected signature of the reliability paradox and indicates that the pooling adopted here rests on the demonstrated group-level equivalence rather than on individual-level stability. Full reproducibility and reliability statistics are provided in the Supplementary Material.

### 3.2 Behavioural results

#### 3.2.1 Session analysis (Session 1 vs. Session 2)

**Figure 2.**
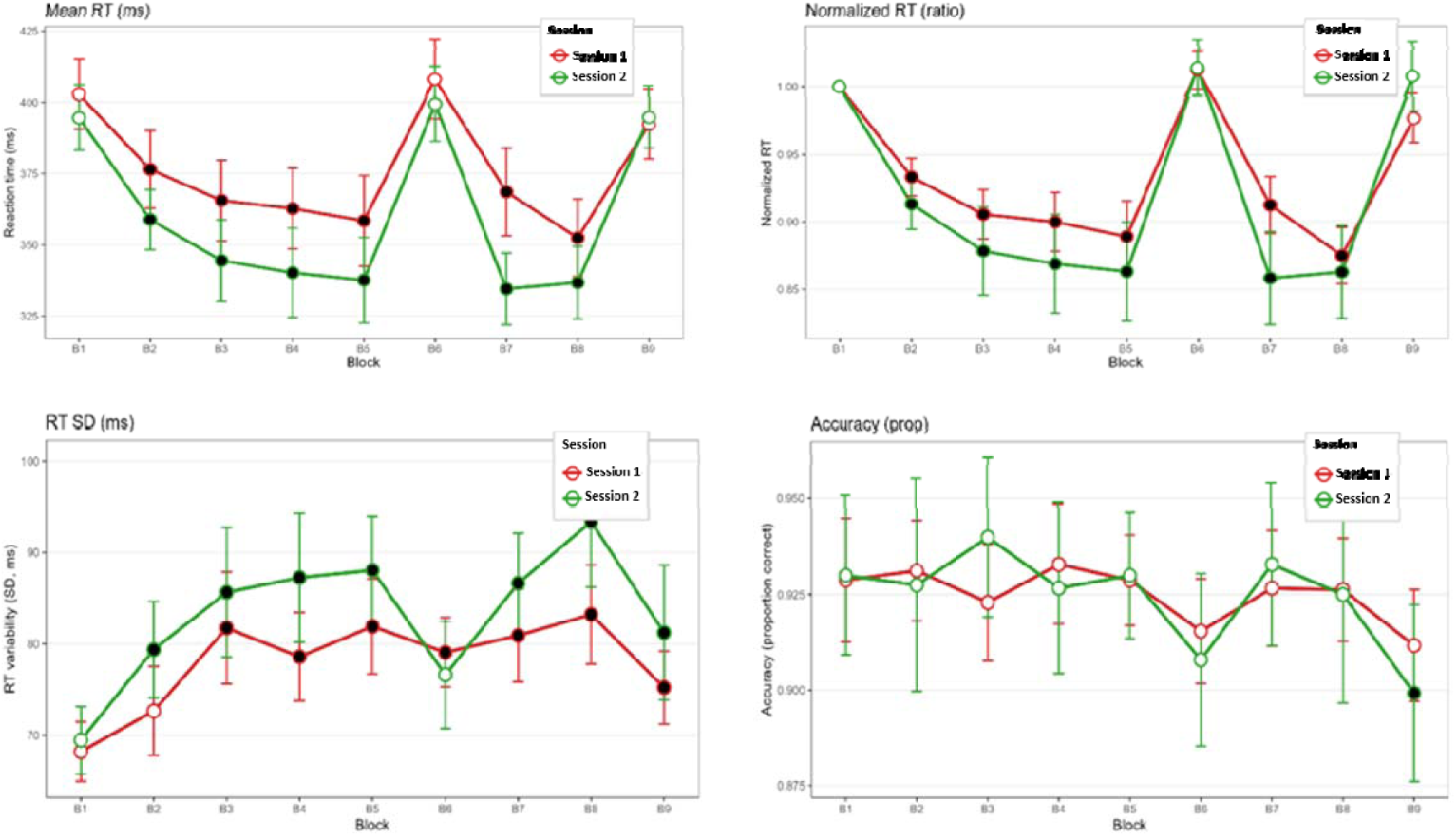
SRTT block-wise behavioural performance for Session 3 (red) and Session 4 (green). Panels: mean absolute RT (upper-left), normalised RT relative to Block 1 (upper-right), RT variability (lower-left), and accuracy (lower-right). Filled black circles indicate blocks differing significantly from Block 1 (paired t-tests, Fisher’s LSD, p < .05). Error bars represent ±1 SEM. n = 20

The ANOVAs showed significant main effects of block in all four behavioural measures, including absolute reaction time, F(2.80, 53.3) = 20.60, ηp^2^ = .520, p < .001; normalised reaction time, F(3.21, 61.0) = 22.89, ηp^2^ = .546, p < .001; error rate, F(3.82, 72.6) = 3.05, ηp^2^ = .138, p = .024; and reaction time variability, F(4.07, 77.4) = 6.33, ηp^2^ = .250, p < .001. In contrast, no significant main effect of Condition was observed for any of these measures, with all p-values greater than .12. The Condition × Block interaction was also not significant, with all p-values greater than .26. This suggests that the pattern of learning across blocks was comparable between the two sessions (Table 2). In addition, the ANOVA conducted for the block-transition analyses of the change scores ΔB5–B6 and ΔB6–B7 did not show significant session differences, with all p-values ≥ .097. This indicates that both the disruption after switching from sequence to random blocks and the recovery after returning to the sequence blocks did not differ across sessions.

**Table 2.** Repeated measures ANOVA for SRTT behavioural measures (Session 1 vs. Session 2).

| Measure | Factor | df, Error | F | $\eta p^2$ | p |
| --- | --- | --- | --- | --- | --- |
| Absolute RT | Condition | 1, 19 | 2.609 | 0.121 | 0.123 |
|  | Block | 2.80, 53.3 | 20.596 | 0.520 | < .001 |
| | Condition × Block | 3.19, 60.6 | 1.360 | 0.067 | 0.263 |
| Normalised RT | Condition | 1, 19 | 0.418 | 0.022 | 0.526 |
|  | Block | 3.21, 61.0 | 22.885 | 0.546 | < .001 |
| | Condition × Block | 3.50, 66.6 | 1.300 | 0.064 | 0.281 |
| Errors | Condition | 1, 19 | 0.002 | 0.000 | 0.965 |
|  | Block | 3.82, 72.6 | 3.048 | 0.138 | 0.024 |
| | Condition × Block | 3.93, 74.6 | 0.761 | 0.039 | 0.552 |
| RT variability | Condition | 1, 19 | 2.078 | 0.099 | 0.166 |
|  | Block | 4.07, 77.4 | 6.330 | 0.250 | < .001 |
| | Condition × Block | 3.57, 67.9 | 0.752 | 0.038 | 0.546 |

Post-hoc comparisons between each block and Block 1 showed significantly shorter reaction times in all sequence blocks, namely B2–B5, B7, and B8, in both sessions. For absolute reaction time, these effects were consistently large, with Cohen’s d values above 0.75. Importantly, the random blocks, B6 and B9, did not differ significantly from Block 1 for either absolute or normalised reaction time in either session. Accuracy remained largely stable across blocks in both sessions. The only exception was Block 9 in Session 4, where participants made significantly more errors than in Block 1, t(19) = 3.561, d = 0.796, p < .01.

#### 3.2.2 Task Version analysis (Task V2 vs. Task V3)

Block again showed a significant within-subject effect for all four behavioural measures (Figure 3). This effect was significant for errors, with all p-values below .025, and significant for the other measures, with all p-values below .001. In contrast, there was no significant main effect of Condition, comparing Task V2 and Task V3, for any measure, with all p-values greater than .47. The Condition × Block interaction was also not significant, with all p-values greater than .17. These findings indicate that the overall learning pattern was comparable between the two task versions (Table 3). Both task versions showed significant reductions in reaction time during all sequence blocks compared with Block 1, while no significant difference was observed between the random blocks. Some differences between the task versions were noted. In Task V2, error rates were significantly higher in the random blocks B6 and B9 compared with Block 1, with p < .01 and p < .001, respectively. This pattern, suggesting an accuracy-related disruption during random blocks, was not observed in Task V3. In contrast, Task V3 showed significantly increased reaction time variability in random Block 6, p < .01, whereas this effect did not reach significance in Task V2, p = .056. However, these version-specific differences did not change the main behavioural pattern of sequence learning at the group level.

**Figure 3.**
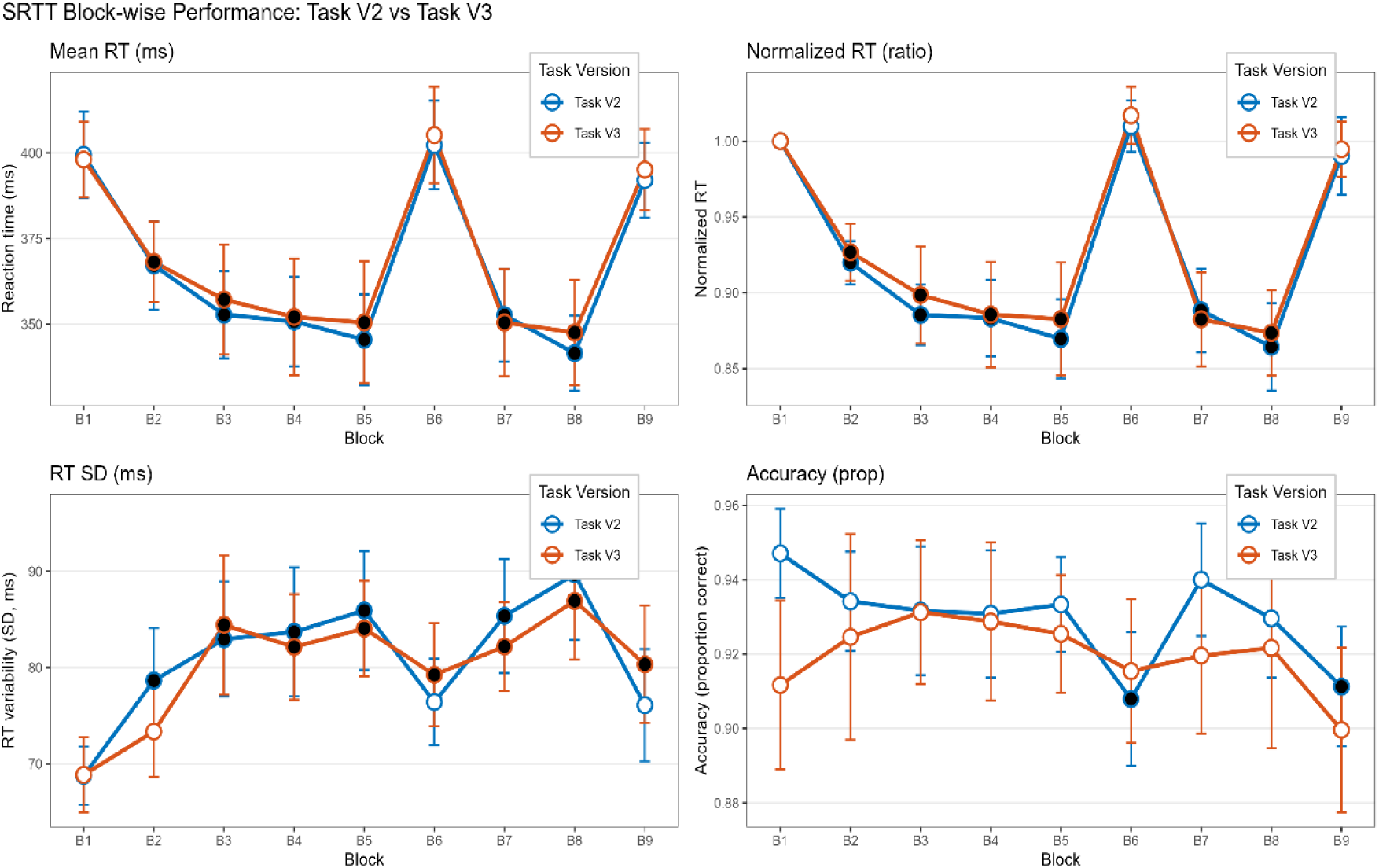
SRTT block-wise behavioural performance for Task V2 (blue) and Task V3 (orange). Panels and conventions as in Figure 2.

**Table 3.** Repeated measures ANOVA for SRTT behavioural measures (Task V2 vs. Task V3).

| Measure | Factor | df, Error | F | $\eta p^2$ | p |
| --- | --- | --- | --- | --- | --- |
| Absolute RT | Condition | 1, 19 | 0.043 | 0.002 | 0.837 |
|  | Block | 2.80, 53.3 | 20.596 | 0.520 | < .001 |
| | Condition × Block | 3.05, 58.0 | 0.091 | 0.005 | 0.967 |
| Normalised RT | Condition | 1, 19 | 0.054 | 0.003 | 0.819 |
|  | Block | 3.21, 61.0 | 22.885 | 0.546 | < .001 |
| | Condition × Block | 3.37, 64.1 | 0.079 | 0.004 | 0.980 |
| RT variability | Condition | 1, 19 | 0.032 | 0.002 | 0.861 |
|  | Block | 4.07, 77.4 | 6.330 | 0.250 | < .001 |
| | Condition × Block | 3.73, 70.8 | 0.482 | 0.025 | 0.736 |
| Errors | Condition | 1, 19 | 0.539 | 0.028 | 0.472 |
|  | Block | 3.82, 72.6 | 3.048 | 0.138 | 0.024 |
| | Condition × Block | 4.14, 78.6 | 1.630 | 0.079 | 0.173 |

### 3.3 Test–Retest Reliability

Test–retest reliability varied depending on block type and the behavioural measure considered (Tables 4 and 5). For absolute reaction time, reliability was good to excellent in the random blocks, including B1, B6, and B9, with ICC (2,1) values of approximately .80 to .85. In contrast, reliability was lower in the sequence blocks, with ICC (2,1) values of approximately .29 to .53 (Figure 4). Accuracy showed generally stronger reliability than reaction time. The highest test–retest stability was observed in the middle sequence blocks, B3 to B5, where ICC (2,1) reached up to .854. Across most blocks, ICC (3,1) values were only slightly higher than ICC (2,1) values. (Tables 4 and 5)

**Table 4.** Intraclass correlation coefficients for reaction time across Sessions 1 and 2.

| Block | Type | ICC (2,1) | 95% CI | ICC(3,1) | 95% CI |
| --- | --- | --- | --- | --- | --- |
| 1 | Random | 0.843 | [0.651, 0.934] | 0.847 | [0.653, 0.936] |
| 2 | Sequence | 0.455 | [0.049, 0.738] | 0.467 | [0.043, 0.749] |
| 3 | Sequence | 0.389 | [−0.032, 0.699] | 0.398 | [−0.042, 0.709] |
| 4 | Sequence | 0.292 | [−0.141, 0.639] | 0.298 | [−0.154, 0.648] |
| 5 | Sequence | 0.414 | [−0.006, 0.715] | 0.421 | [−0.015, 0.722] |
| 6 | Random | 0.827 | [0.619, 0.927] | 0.828 | [0.615, 0.928] |
| 7 | Sequence | 0.346 | [−0.051, 0.665] | 0.386 | [−0.056, 0.702] |
| 8 | Sequence | 0.532 | [0.144, 0.782] | 0.539 | [0.138, 0.788] |
| 9 | Random | 0.803 | [0.566, 0.917] | 0.796 | [0.553, 0.914] |

**Table 5.** Intraclass correlation coefficients for accuracy across Sessions 1 and 2.

| Block | Type | ICC (2,1) | 95% CI | ICC(3,1) | 95% CI |
| --- | --- | --- | --- | --- | --- |
| 1 | Random | 0.473 | [0.036, 0.754] | 0.461 | [0.035, 0.745] |
| 2 | Sequence | 0.541 | [0.131, 0.791] | 0.529 | [0.125, 0.783] |
| 3 | Sequence | 0.807 | [0.579, 0.919] | 0.817 | [0.595, 0.924] |
| 4 | Sequence | 0.854 | [0.670, 0.940] | 0.850 | [0.660, 0.938] |
| 5 | Sequence | 0.696 | [0.370, 0.868] | 0.685 | [0.358, 0.862] |
| 6 | Random | 0.684 | [0.354, 0.862] | 0.675 | [0.342, 0.857] |
| 7 | Sequence | 0.464 | [0.027, 0.749] | 0.453 | [0.025, 0.741] |
| 8 | Sequence | 0.586 | [0.195, 0.814] | 0.574 | [0.187, 0.806] |
| 9 | Random | 0.706 | [0.399, 0.872] | 0.703 | [0.389, 0.871] |

**Figure 4.**
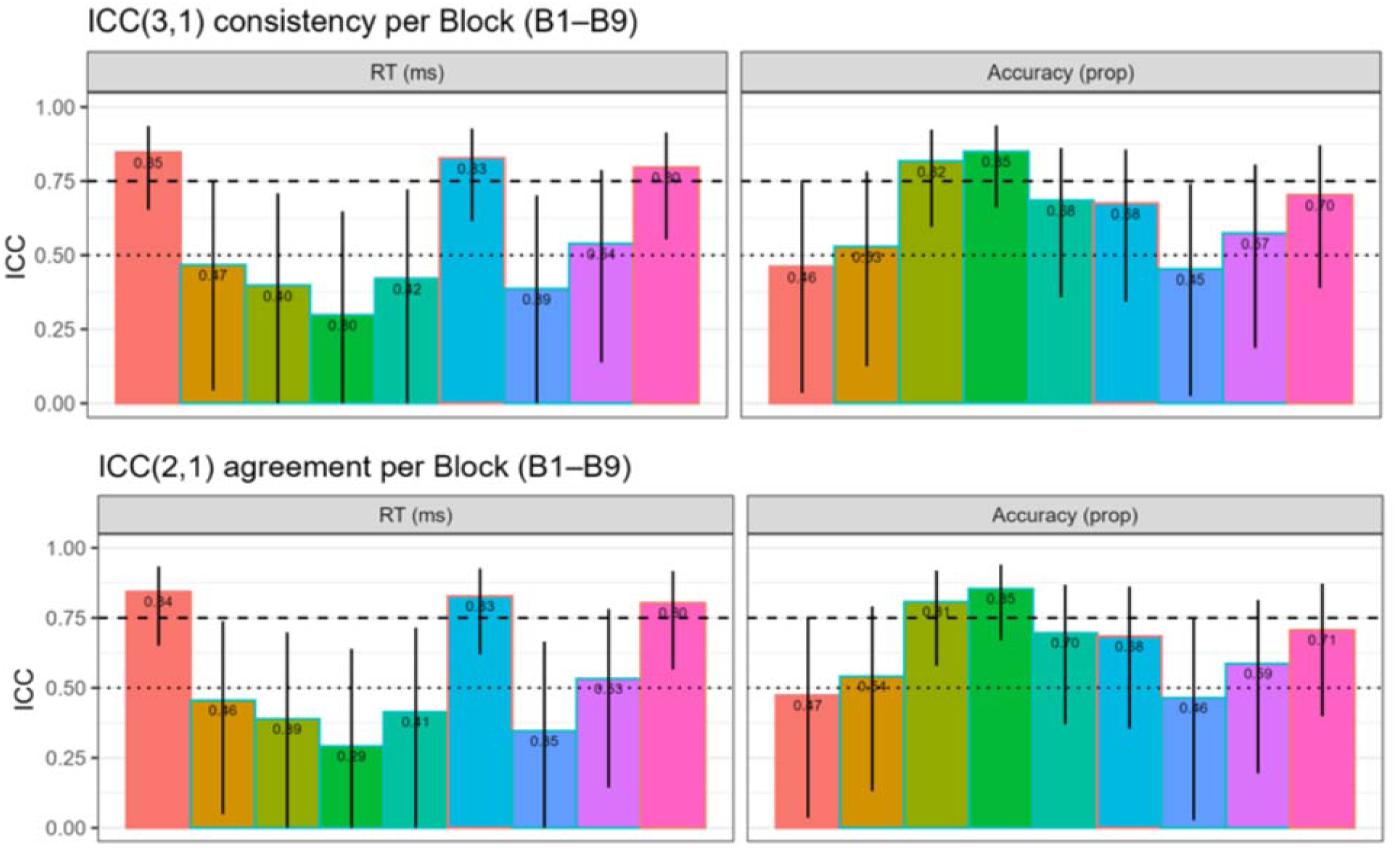
Test–retest reliability of SRTT performance across Sessions 3 and 4: ICC(3,1) consistency (top panels) and ICC(2,1) absolute agreement (bottom panels) for reaction time (left) and accuracy (right). Bars show block-wise intraclass correlation coefficients (B1–B9), with random blocks (B1, B6, B9) in distinct colour. Error bars: 95% bootstrap confidence intervals. Dashed and dotted lines indicate good (ICC ≥ .75) and moderate (ICC ≥ .50) thresholds.

### 3.4 Whole-brain BOLD activation

Whole-brain activation was examined first for the task blocks relative to rest (SEQ > rest (Baseline) and RND > rest (Baseline)), and then for the direct contrast between sequence and random blocks (SEQ > RND).

For the SEQ > Baseline contrast, four clusters survived cluster-level FWE correction (k = 20,204 / 1,182 / 297 / 177). The dominant cluster included the right middle cingulate and paracingulate gyrus (12, 8, 44; T = 11.53) and several distinct anatomical sub-peaks. These included the bilateral supplementary motor area (SMA; e.g., −2, −2, 56; T = 8.42), left-lateralised primary motor and somatosensory cortex (left precentral gyrus, −36, −20, 58, T = 9.45; left postcentral gyrus, −40, −28, 60, T = 10.02), right sensorimotor and opercular cortex (right postcentral gyrus, 62, −12, 30, T = 8.76; right central operculum, 38, −8, 20, T = 8.16), the left superior parietal lobule (−36, −42, 54; T = 7.96), and superior frontal/dorsal premotor cortex (−16, −6, 70; T = 9.38). Three further clusters were located in the cerebellum, and included the right lobule VI with vermis IV/V (k = 1,182; peak T = 7.66), the right lobule VIII (k = 297; peak T = 6.78), and the left lobule VI (k = 177; peak T = 6.92) respectively.

For the RND > Baseline contrast, three clusters survived correction (k = 17,873 / 1,182 / 422). This contrast largely reproduced cluster I of the SEQ pattern, with comparable peak intensities in the SMA (−2, −2, 56; T = 9.03) and in bilateral primary motor and somatosensory cortex (right postcentral gyrus, 62, −12, 30, T = 11.54; left postcentral gyrus, −40, −28, 56, T = 10.62), as well as the clusters including cerebellar lobules VI and VIII. Because both block types impose the same overt visuomotor response demand, the SEQ > Baseline and RND > Baseline contrasts cannot disentangle the common motor-execution network from sequence-specific network components. Sequence-specific effects were therefore isolated using the direct SEQ > RND contrast.

#### Sequence-specific activation (SEQ > RND)

Six clusters survived cluster-level FWE correction for the SEQ > RND contrast (Table 5). The largest cluster was mainly located in the left superior parietal lobule and extended into neighboring sensorimotor and dorsal premotor regions (k = 1,518; peak T = 5.81) (Figure 5). A second large cluster included the left lingual gyrus, fusiform gyrus, lateral occipital cortex, and left cerebellar Crus I (k = 568; peak T = 5.84). Further significant clusters were found in bilateral parietal and Rolandic opercular regions, consistent with activation of the secondary somatosensory cortex (S2). These included a left-sided cluster extending into the insular cortex (k = 475; peak T = 6.38) and a right-sided cluster in the corresponding opercular region (k = 260; peak T = 5.00). Two additional clusters were located in medial premotor regions. One cluster involved the left supplementary motor area (SMA) and paracingulate cortex (k = 251; peak T = 5.55), while the other involved the right SMA and middle cingulate cortex (k = 195; peak T = 5.60).

**Figure 5.**
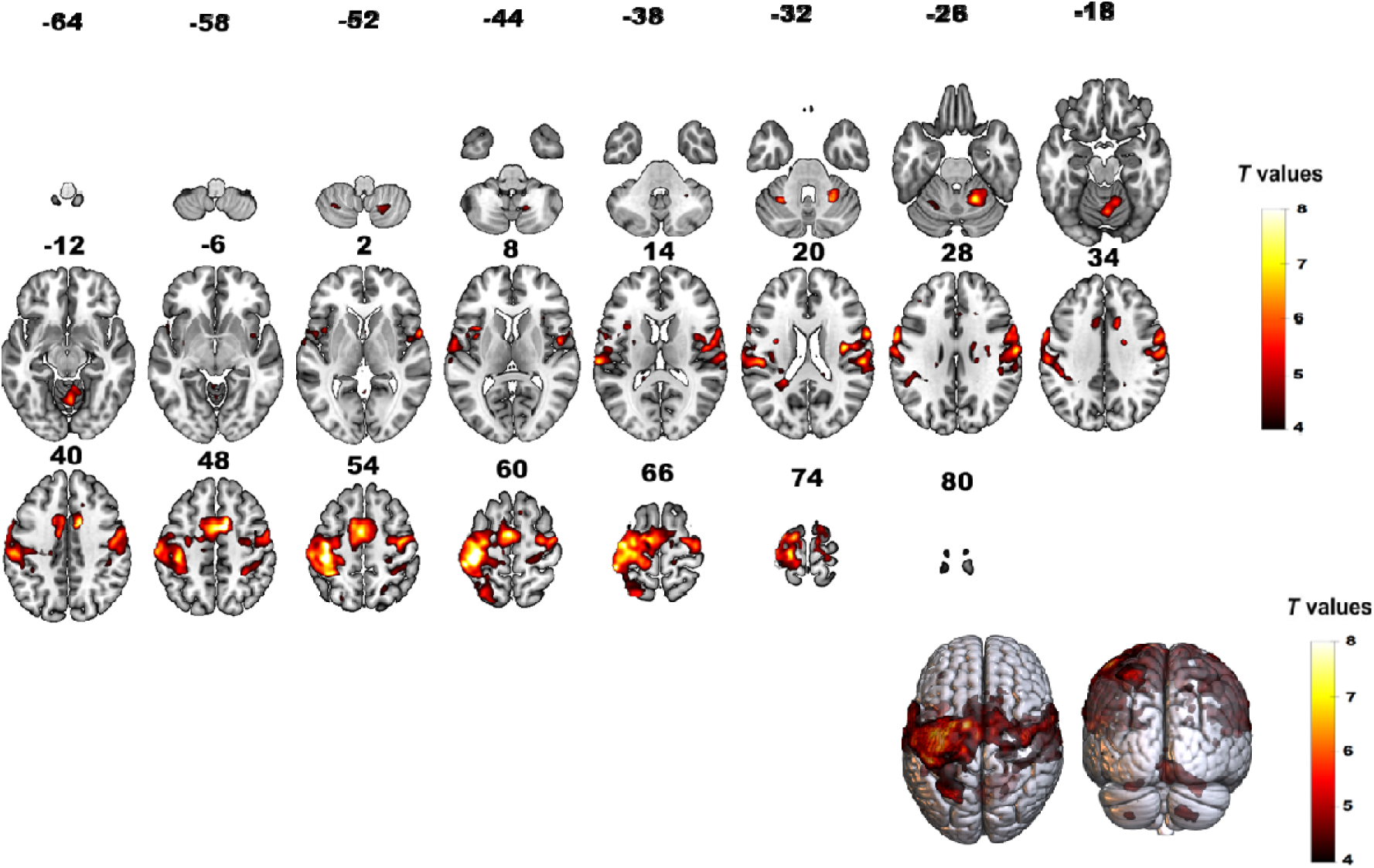
**(a).** Whole-brain activation map for the contrast SEQ > Baseline, thresholded at cluster-level family-wise error correction (FWEc) p < 0.05, with a voxel-wise cluster-forming threshold of p < 0.001 uncorrected. Activations are overlaid on the MNI152 T1 template shown as axial slices (Z-coordinates in mm, top) and as dorsal and posterior 3D cortical surface renderings (bottom right). Warm colours represent positive T-values. Cluster sizes, peak T-values, peak MNI coordinates, and anatomical labels are reported in Table **6(a).**

**Table 6(a).**
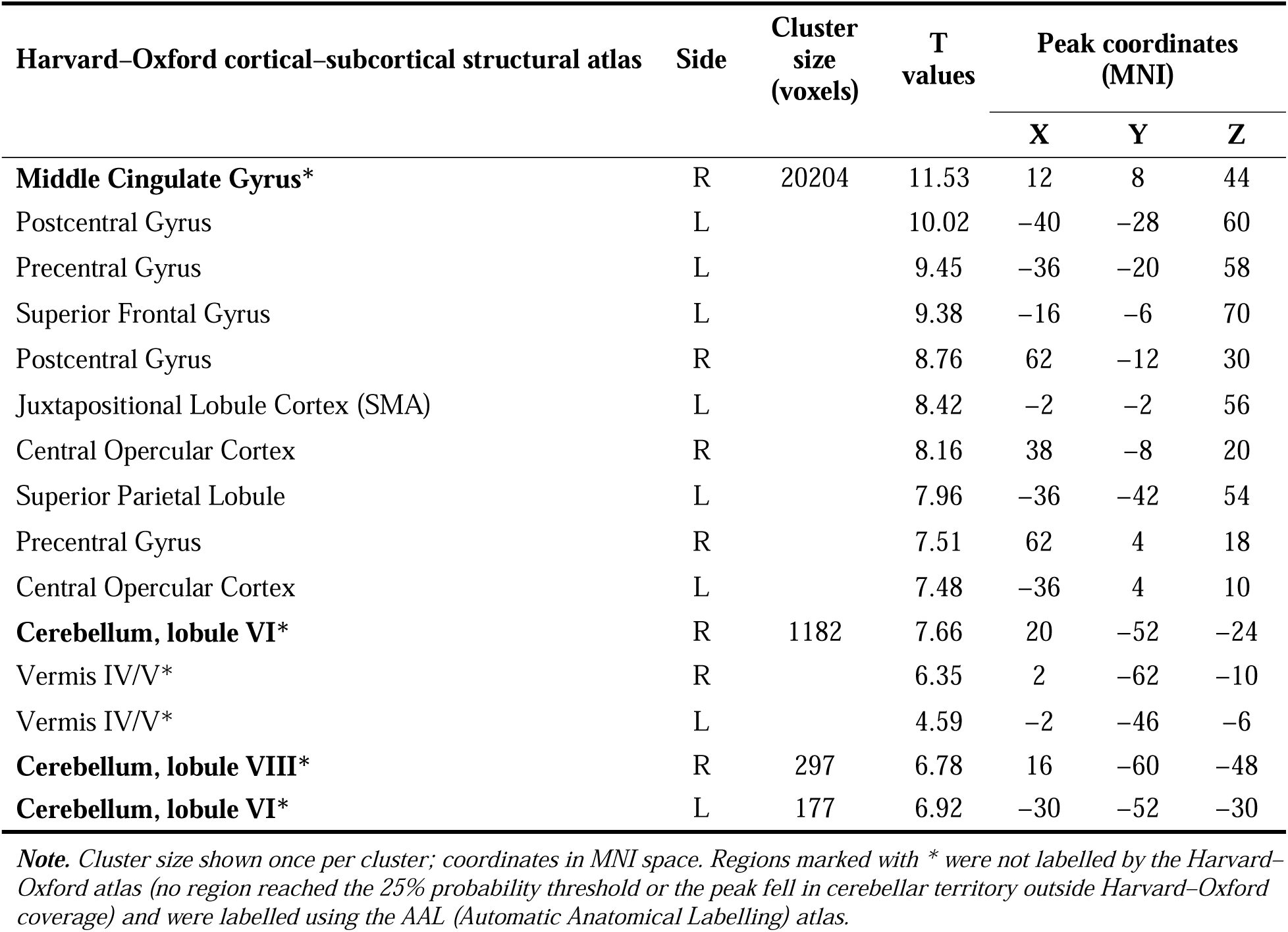
Significant brain activation for the contrast SEQ > Baseline.

| Harvard–Oxford cortical–subcortical structural atlas | Side | Cluster size (voxels) | T values | Peak coordinates (MNI) |  |  |
| --- | --- | --- | --- | --- | --- | --- |
|  |  |  |  | X | Y | Z |
| **Middle Cingulate Gyrus\*** | R | 20204 | 11.53 | 12 | 8 | 44 |
| Postcentral Gyrus | L |  | 10.02 | -40 | -28 | 60 |
| Precentral Gyrus | L |  | 9.45 | -36 | -20 | 58 |
| Superior Frontal Gyrus | L |  | 9.38 | -16 | -6 | 70 |
| Postcentral Gyrus | R |  | 8.76 | 62 | -12 | 30 |
| Juxtapositional Lobule Cortex (SMA) | L |  | 8.42 | -2 | -2 | 56 |
| Central Opercular Cortex | R |  | 8.16 | 38 | -8 | 20 |
| Superior Parietal Lobule | L |  | 7.96 | -36 | -42 | 54 |
| Precentral Gyrus | R |  | 7.51 | 62 | 4 | 18 |
| Central Opercular Cortex | L |  | 7.48 | -36 | 4 | 10 |
| **Cerebellum, lobule VI\*** | R | 1182 | 7.66 | 20 | -52 | -24 |
| Vermis IV/V* | R |  | 6.35 | 2 | -62 | -10 |
| Vermis IV/V* | L |  | 4.59 | -2 | -46 | -6 |
| **Cerebellum, lobule VIII\*** | R | 297 | 6.78 | 16 | -60 | -48 |
| **Cerebellum, lobule VI\*** | L | 177 | 6.92 | -30 | -52 | -30 |

*Note.* Cluster size shown once per cluster; coordinates in MNI space. Regions marked with \* were not labelled by the Harvard–Oxford atlas (no region reached the 25% probability threshold or the peak fell in cerebellar territory outside Harvard–Oxford coverage) and were labelled using the AAL (Automatic Anatomical Labelling) atlas.

**Figure 5.**
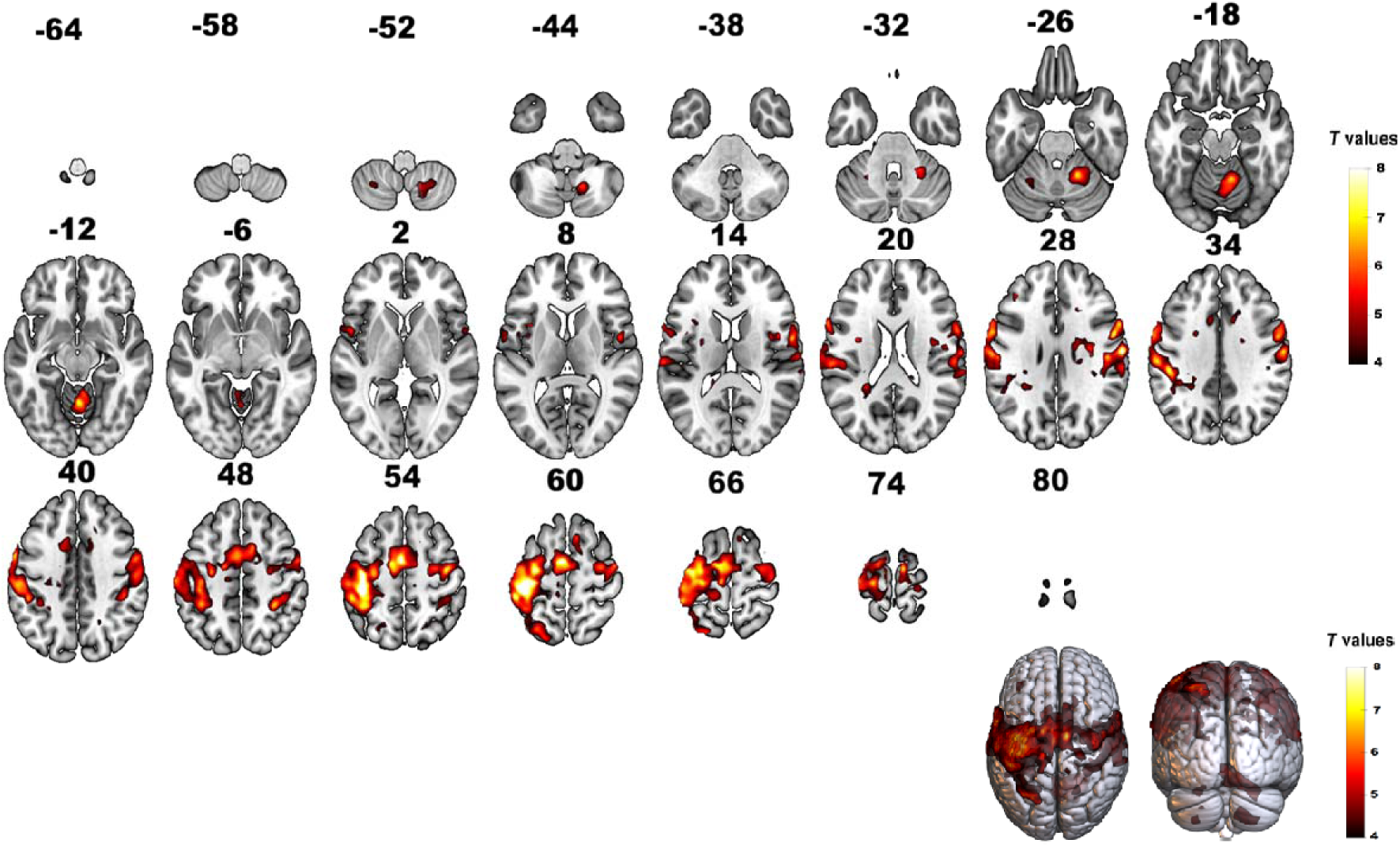
**(b).** Whole-brain activation map for the contrast RND > Baseline, thresholded at cluster-level family-wise error correction (FWEc) p < 0.05, with a voxel-wise cluster-forming threshold of p < 0.001 uncorrected. Activations are overlaid on the MNI152 T1 template shown as axial slices (Z-coordinates in mm, top) and as dorsal and posterior 3D cortical surface renderings (bottom right). Warm colours represent positive T-values. Cluster sizes, peak T-values, peak MNI coordinates, and anatomical labels are reported in Table 6(b).

**Figure 5.**
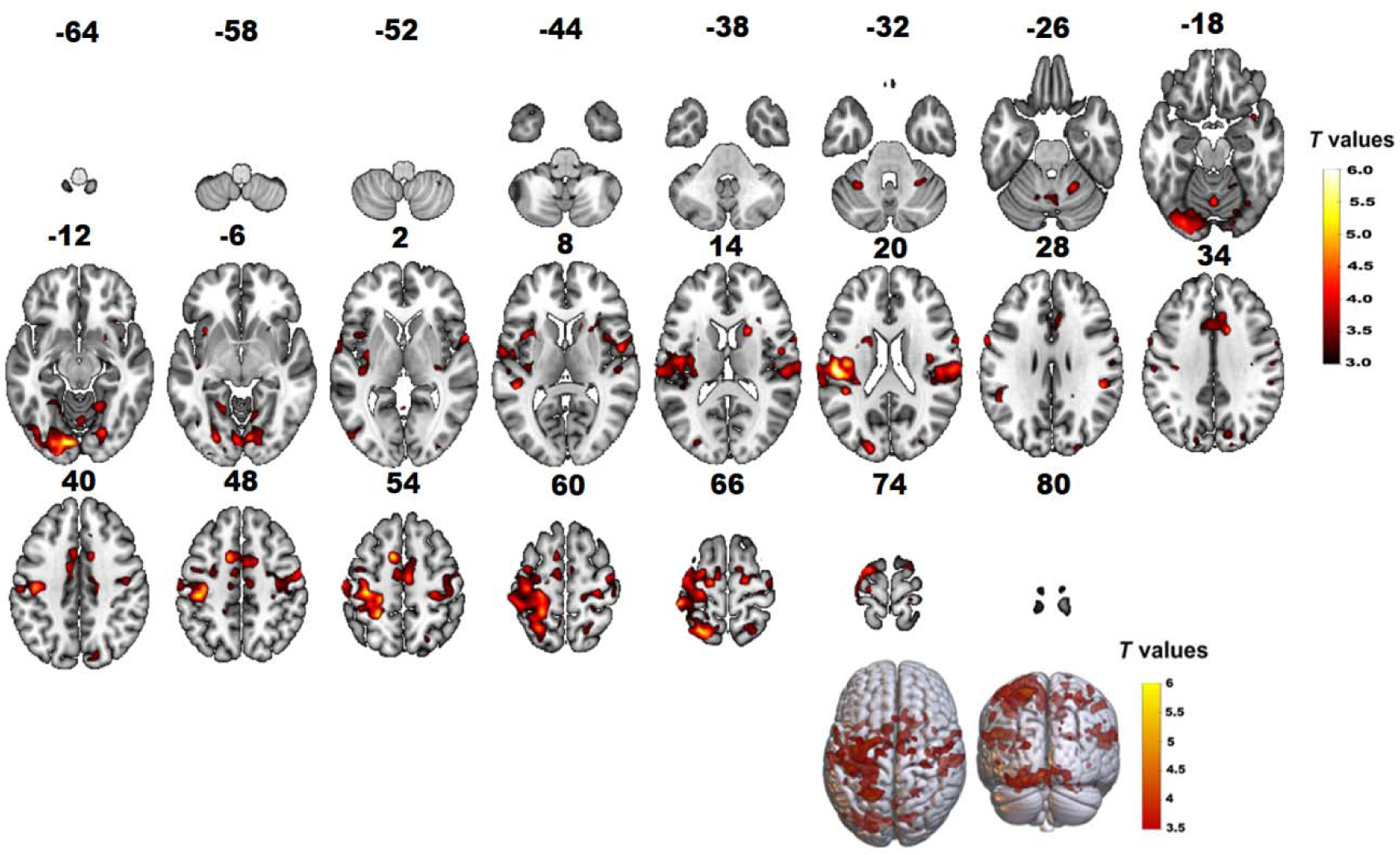
**(c).** Whole-brain activation map for the contrast SEQ > RND, thresholded at cluster-level family-wise error correction (FWEc) p < 0.05, with a voxel-wise cluster-forming threshold of p < 0.001 uncorrected. Activations are overlaid on the MNI152 T1 template shown as axial slices (Z-coordinates in mm, top) and as dorsal and posterior 3D cortical surface renderings (bottom right). Warm colours represent positive T-values. Cluster sizes, peak T-values, peak MNI coordinates, and anatomical labels are reported in Table 6(c).

**Table 6(b).** Significant brain activation for the contrast RND > Baseline.

| Harvard–Oxford cortical–subcortical structural atlas | Side | Cluster size (voxels) | T values | Peak coordinates (MNI) |  |  |
| --- | --- | --- | --- | --- | --- | --- |
|  |  |  |  | X | Y | Z |
| **Postcentral Gyrus** | R | 17873 | 11.54 | 62 | -12 | 30 |
| Postcentral Gyrus | L |  | 10.62 | -40 | -28 | 56 |
| Supramarginal Gyrus* | R |  | 9.61 | 48 | -18 | 26 |
| Juxtapositional Lobule Cortex (SMA) | L |  | 9.03 | -2 | -2 | 56 |
| Precentral Gyrus | R |  | 8.56 | 54 | 2 | 30 |
|  |  |  |  | X | Y | Z |
| Precentral Gyrus | L |  | 8.43 | −38 | −14 | 54 |
| Supplementary Motor Area* | L |  | 8.18 | −12 | −6 | 52 |
| Superior Frontal Gyrus | R |  | 7.97 | 10 | −10 | 74 |
| Superior Frontal Gyrus | L |  | 7.72 | −10 | −8 | 68 |
| Superior Parietal Lobule | L |  | 7.71 | −38 | −40 | 52 |
| **Cerebellum, lobule VI\*** | R | 1182 | 7.32 | 20 | −52 | −24 |
| Vermis IV/V* | R |  | 7.30 | 2 | −56 | −12 |
| Cerebellum, lobule IV/V* | R |  | 7.22 | 10 | −54 | −16 |
| Cerebellum, lobule IV/V* | L |  | 5.08 | −4 | −48 | −6 |
| Vermis IV/V* | M |  | 3.91 | 0 | −52 | 0 |
| **Cerebellum, lobule VIII\*** | R | 422 | 6.56 | 16 | −60 | −48 |
| Vermis VIII* | R |  | 4.11 | 4 | −70 | −36 |

**Table 6(c).** Significant brain activation for the contrast SEQ > RND.

| Harvard–Oxford cortical–subcortical structural atlas | Side | Cluster size (voxels) | T values | Peak coordinates (MNI) |  |  |
| --- | --- | --- | --- | --- | --- | --- |
|  |  |  |  | X | Y | Z |
| **Lateral Occipital Cortex superior division** | L | 1518 | 5.81 | −18 | −58 | 68 |
| Postcentral Gyrus* | L |  | 5.77 | −24 | −42 | 54 |
| Precentral Gyrus | L |  | 5.50 | −34 | −24 | 54 |
| Superior Parietal Lobule | L |  | 5.48 | −26 | −56 | 66 |
| Superior Frontal Gyrus | L |  | 5.18 | −24 | −8 | 70 |
| Middle Frontal Gyrus | L |  | 4.16 | −36 | −4 | 58 |
| **Lingual Gyrus** | L | 568 | 5.84 | −10 | −84 | −12 |
|  |  |  |  | X | Y | Z |
| Occipital Fusiform Gyrus | L |  | 4.82 | −24 | −84 | −12 |
| Cerebellum, Crus I* | L |  | 4.09 | −36 | −84 | −20 |
| Lateral Occipital Cortex inferior division | L |  | 4.05 | −40 | −84 | −12 |
| **Central Opercular Cortex** | L | 475 | 6.38 | −50 | −22 | 20 |
| Parietal Operculum Cortex | L |  | 4.62 | −62 | −24 | 16 |
| Insular Cortex | L |  | 4.33 | −34 | −22 | 18 |
| Rolandic Operculum* | L |  | 4.02 | −30 | −32 | 20 |
| **Parietal Operculum Cortex** | R | 260 | 5.00 | 56 | −22 | 18 |
| **Supplementary Motor Area\*** | L | 251 | 5.55 | −10 | 8 | 54 |
| Juxtapositional Lobule Cortex (SMA) | L |  | 5.21 | −12 | 6 | 46 |
| Paracingulate Gyrus | L |  | 4.28 | −6 | 10 | 42 |
| **Middle Cingulate Gyrus\*** | R | 195 | 5.60 | 14 | 14 | 32 |
| Juxtapositional Lobule Cortex (SMA) | R |  | 5.06 | 6 | −10 | 56 |
| Supplementary Motor Area* | R |  | 4.23 | 14 | 2 | 50 |
| Paracingulate Gyrus | R |  | 3.97 | 10 | 26 | 34 |
| Precentral Gyrus | R |  | 3.70 | 10 | −16 | 46 |

Stability of SEQ > RND activation across sessions and task versions: We a examined whether the SEQ > RND activation pattern was stable across repeated measurements and task versions. Paired-samples t-tests were performed on the SEQ > RND contrast images, comparing Session 1 with Session 2 and Task V2 with Task V3. The same statistical threshold as in the main SEQ > RND analysis was applied.

No clusters survived cluster-level FWE correction at p < .05 in any of these comparisons. For the Session 1 > Session 2 contrast, no clusters were observed even at the uncorrected threshold. In the remaining contrasts, only a small number of low-extent clusters appeared at p < .001 uncorrected, with peak T-values ≤ 5.14 and cluster sizes ≤ 35 voxels. However, none of these clusters survived correction for multiple comparisons, with all cluster-level p-FWE values ≥ .762 and all peak-level p-FWE values ≥ .158.

The reverse contrast (RND > SEQ) yielded no significant activation at the whole-brain FWE-corrected threshold (p < .05). At an uncorrected threshold of p < .001 only a single small periventricular cluster emerged (peak t = 5.91), which was not anatomically interpretable and did not survive correction.

### 3.5 Seed-based connectivity (SBC) of the motor network

Before describing the connectivity effects, order (session) and task-version effects were first examined and did not differ at the group level (only 2 of 50 edge-tests differed between sessions, and none between versions, after correction). The the full reproducibility and reliability statistics, and the basis for relying on the group-level t-tests, are reported in Section 3.1 and the Supplementary Material.

Across all five task conditions (the five task phases, each modelled separately: Initial RND, Early SEQ, Middle SEQ, Late SEQ, and Interleaved RND), the predefined five-region motor network formed one significant connectivity cluster (Figure 6) in the omnibus F-test, with all cluster-level p-FDR values below .001 (Table 7). Within each block condition, seven to nine of the ten possible pairwise connections survived within-cluster FWE correction. The overall strength of the network varied across the five conditions in a non-linear pattern. It was already high during the initial random baseline block (F = 58.77), decreased during early sequence exposure (Early SEQ: F = 34.76), increased again during the middle sequence phase (Middle SEQ: F = 60.42), became more moderate during the late sequence phase (Late SEQ: F = 41.38), and was strongest during the interleaved random blocks (F = 72.55)

**Figure 6.**
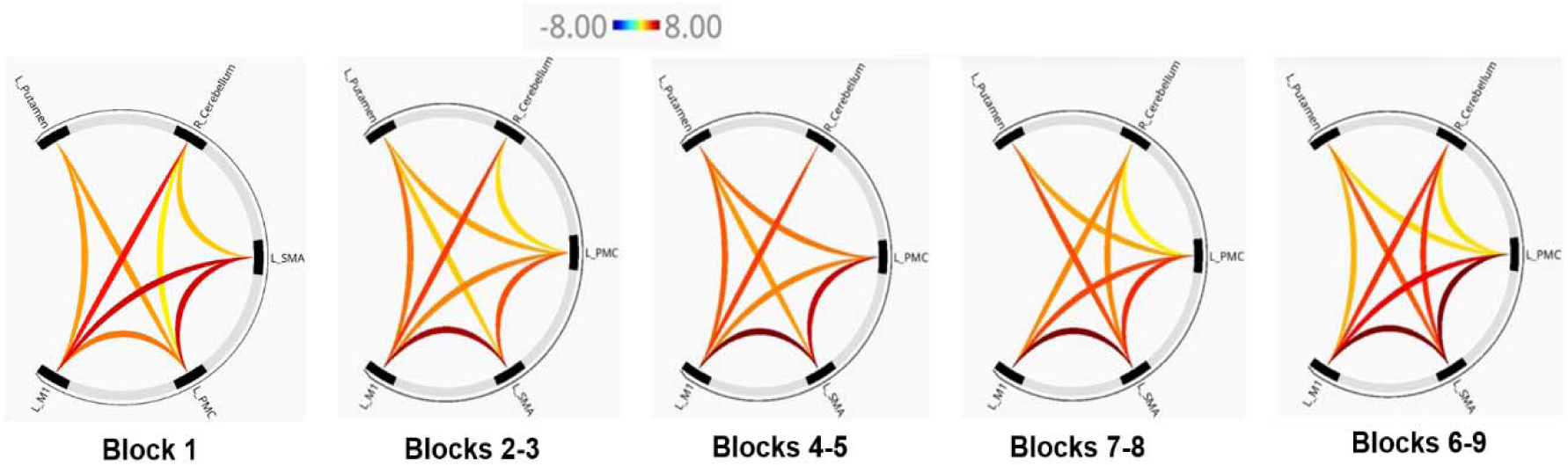
Seed-based functional connectivity. CONN toolbox connectome ring plots of the motor network across the five task conditions are shown. Each panel corresponds to one condition: RND (Block 1), Early SEQ (Blocks 2–3), Middle SEQ (Blocks 4–5), Late SEQ (Blocks 7–8), and Interleaved RND (Blocks 6, 9). Each panel displays the same five ROIs — L_Putamen, R_Cerebellum, and the three left-hemispheric cortical motor nodes (L_M1, L_SMA, L_PMC). Each arc represents a significant within-cluster connection; arc color encodes the T statistic on a blue–white–red scale from −8.00 to +8.00 (dark red = strongest positive coupling).

**Table 7.** Cluster-level omnibus statistics for each of the five SBC conditions.

| Condition (blocks) | Omnibus F | p-unc | p-FDR | Sig. connections |
| --- | --- | --- | --- | --- |
| Initial RND (Block 1) | 58.77 | < 0.001 | < 0.001 | 8 |
| Early SEQ (Blocks 2–3) | 34.76 | < 0.001 | < 0.001 | 8 |
| Middle SEQ (Blocks 4–5) | 60.42 | < 0.001 | < 0.001 | 7 |
| Late SEQ (Blocks 7–8) | 41.38 | < 0.001 | < 0.001 | 8 |
| Interleaved RND (Blocks 6,9) | 72.55 | < 0.001 | < 0.001 | 9 |

**Figure 7.**
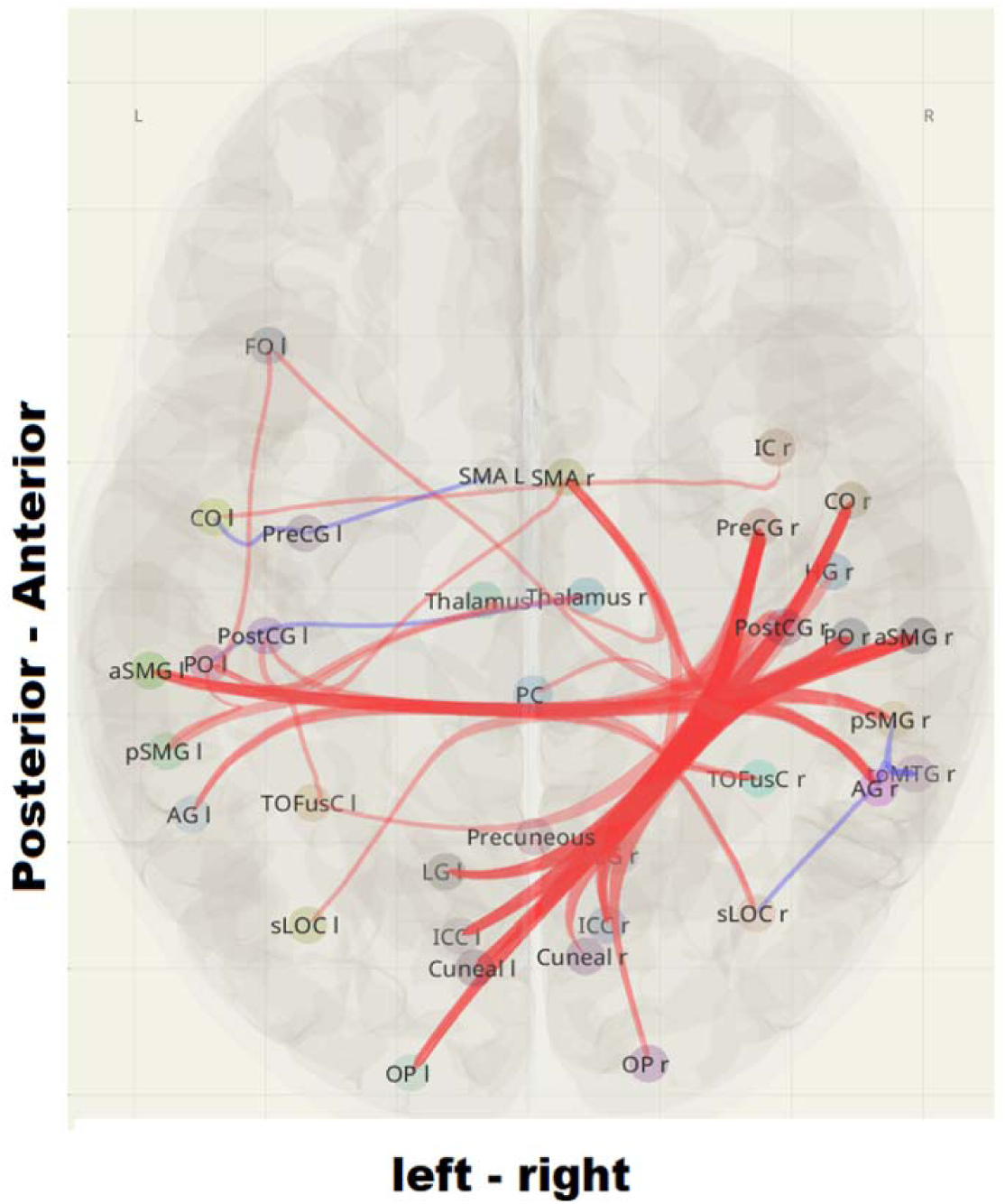
General Psychophysiological interaction (gPPI). The axial glass-brain projection shows the significant task-modulated functional connections for the SEQ > RND contrast for the combined blocks of each condition. Nodes are positioned at the MNI coordinates of each ROI and labelled with Harvard–Oxford atlas abbreviations. Red edges indicate increased coupling during SEQ relative to RND; blue edges indicate decreased coupling. Edge thickness reflects the magnitude of the T-statistic. Axes: X = left–right; Y = posterior–anterior (MNI mm). L = left; R = right.

The connection between L_M1 and L_SMA was the strongest, or among the strongest, in every condition. Its strength increased from T = 6.66 in the Initial RND condition to T = 8.48 in the Interleaved RND condition. It was thus the most consistent pairwise coupling in the network across all conditions (Table 8). Two additional connections were especially relevant for interpreting the learning-related changes. First, the L_SMA–L_Putamen connection was not significant during the initial random baseline but became stronger across the sequence-learning phases. Its T-values increased from non-significant at baseline to 2.82 in Early SEQ, 3.62 in Middle SEQ, 4.83 in Late SEQ, and 4.65 in Interleaved RND. This pattern suggests that increasing SMA–putamen Second, the L_SMA–R_Cerebellum connection was weakly significant at baseline (T = 2.89), was not significant during Early and Middle SEQ, and became significant again during Late SEQ and Interleaved RND, with T-values of 3.89 and 5.38, respectively.

**Table 8.**
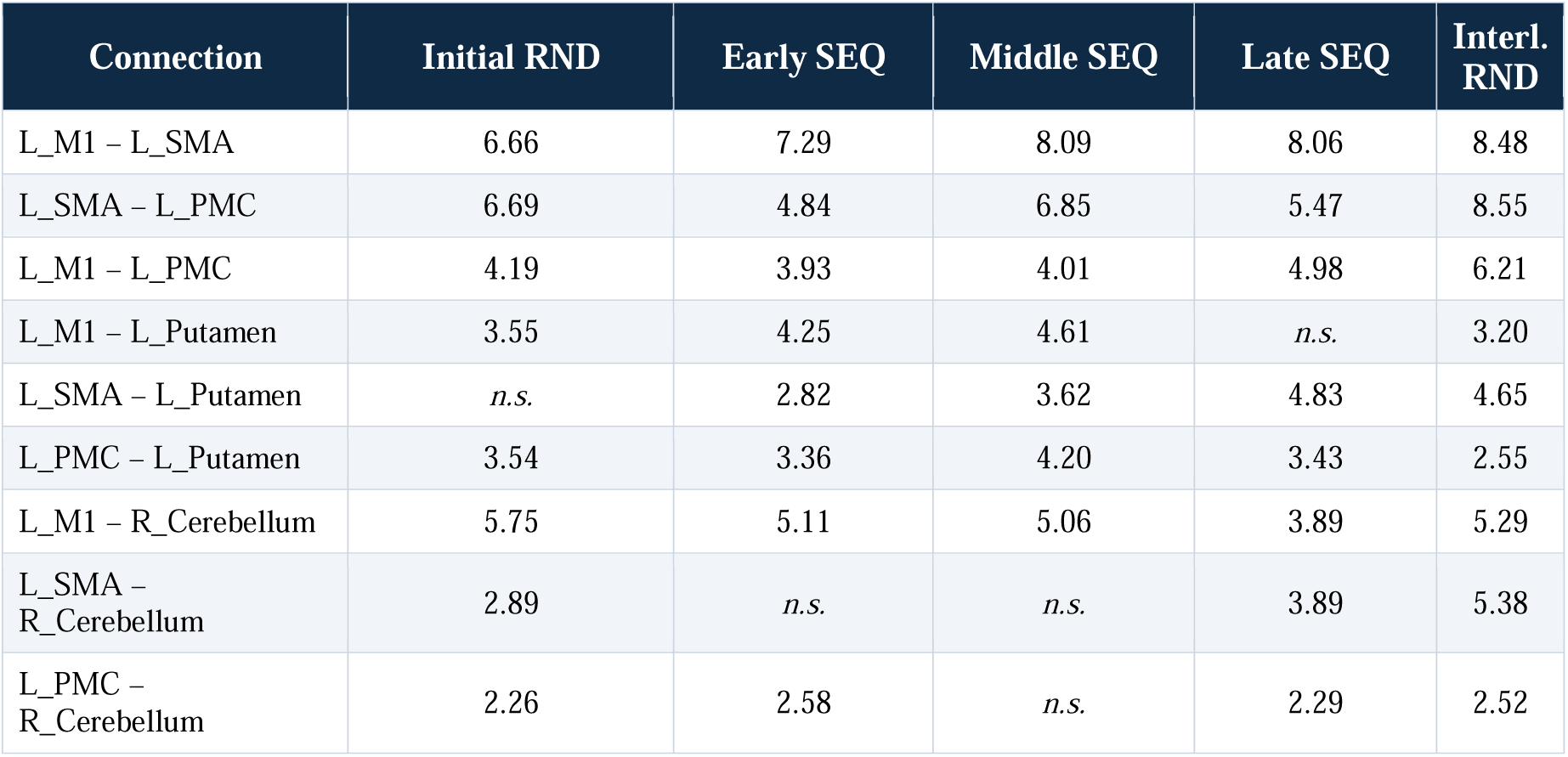
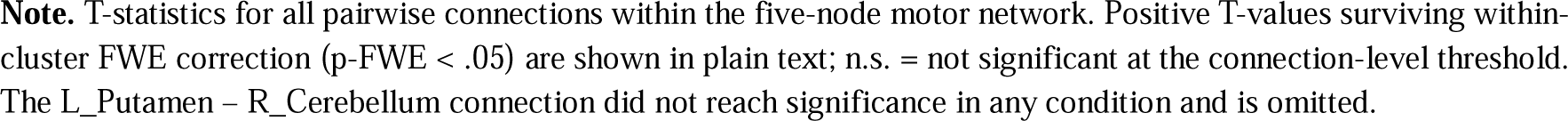
Connection-level T statistics across the five task conditions.

| Connection | Initial RND | Early SEQ | Middle SEQ | Late SEQ | Interl. RND |
| --- | --- | --- | --- | --- | --- |
| L_M1 – L_SMA | 6.66 | 7.29 | 8.09 | 8.06 | 8.48 |
| L_SMA – L_PMC | 6.69 | 4.84 | 6.85 | 5.47 | 8.55 |
| L_M1 – L_PMC | 4.19 | 3.93 | 4.01 | 4.98 | 6.21 |
| L_M1 – L_Putamen | 3.55 | 4.25 | 4.61 | *n.s.* | 3.20 |
| L_SMA – L_Putamen | *n.s.* | 2.82 | 3.62 | 4.83 | 4.65 |
| L_PMC – L_Putamen | 3.54 | 3.36 | 4.20 | 3.43 | 2.55 |
| L_M1 – R_Cerebellum | 5.75 | 5.11 | 5.06 | 3.89 | 5.29 |
| L_SMA – R_Cerebellum | 2.89 | *n.s.* | *n.s.* | 3.89 | 5.38 |
| L_PMC – R_Cerebellum | 2.26 | 2.58 | *n.s.* | 2.29 | 2.52 |

**Note.** T-statistics for all pairwise connections within the five-node motor network. Positive T-values surviving within-cluster FWE correction ( $p\text{-FWE} < .05$ ) are shown in plain text; n.s. = not significant at the connection-level threshold. The L\_Putamen – R\_Cerebellum connection did not reach significance in any condition and is omitted.

**Table 9.** Functional network clusters surviving cluster-level FDR correction (SEQ > RND combined Sessions 1–2).

| Cluster | F | p-unc | p-FDR | Network label |
| --- | --- | --- | --- | --- |
| 1 | 15.14 | 0.000168 | 0.020 | Right-lateralised visuomotor integration network |
| 2 | 11.36 | 0.000737 | 0.039 | Bilateral parieto-premotor–thalamic network |
| 3 | 10.74 | 0.000963 | 0.039 | Left sensorimotor decoupling (automatisation) |
| 4 | 9.72 | 0.001535 | 0.046 | Somatosensory–ventral visual integration |

### 3.6 Generalised psychophysiological interaction (gPPI) for SEQ > RND

Before describing the sequence-specific connectivity effects, order (session) and task-version effects were examined and did not differ at the group level. The exploratory gPPI SEQ > RND pattern did not differ between sessions or versions, with full details reported in Section 3.1 and the Supplementary Material.

The gPPI analysis identified four functional network clusters in which connectivity was significantly modulated by sequence learning in the SEQ > RND contrast (Table 8)

#### Cluster 1: Right lateralised visuomotor integration

The largest and strongest cluster showed increased connectivity within a broad visuomotor network (F = 15.14, p-FDR = .020). This network included bilateral visual regions, such as the lingual gyrus, intracalcarine cortex, cuneal cortex, and occipital pole, together with right-sided sensorimotor and parietal areas. These included the right precentral gyrus, right central opercular cortex, right parietal operculum, right anterior supramarginal gyrus, and precuneus.

The strongest connections that survived within-cluster FWE correction were found between the right lingual gyrus and the right central opercular cortex (T = 5.49, p-FWE = .001), between the right lingual gyrus and the right precentral gyrus (T = 5.26, p-FWE = .001), and between the precuneus and the right precentral gyrus (T = 5.16, p-FWE = .009). Taken together, these connections indicate stronger visual–sensorimotor coupling during sequence blocks than during random blocks.

#### Cluster 2: Bilateral parieto-premotor–thalamic network

The second cluster showed significant sequence-related modulation in a bilateral parieto-premotor and thalamic network (F = 11.36, p-FDR = .039). This cluster included subdivisions of the bilateral supramarginal gyrus and showed increased connectivity with the right SMA, left parietal operculum, right angular gyrus, bilateral thalamus, and right superior lateral occipital cortex during sequence blocks.

This cluster also included some connections with reduced connectivity during SEQ compared with RND. These decreases were observed between the right temporo-occipital middle temporal gyrus and the right posterior supramarginal gyrus and angular gyrus (T = −2.57 and −2.52), as well as between the right thalamus and the left parietal operculum (T = −2.66).

#### Cluster 3: Left sensorimotor decoupling

The third cluster was smaller and mainly located in the left hemisphere (F = 10.74, p-FDR = .039). It was characterised mainly by reduced connectivity during SEQ relative to RND. Specifically, the left SMA showed weaker coupling with the left precentral gyrus (T = −3.00), and the left central opercular cortex showed weaker coupling with the left precentral gyrus (T = −2.37). Taken together, these effects indicate reduced left premotor–primary motor coupling during sequence relative to random blocks.

#### Cluster 4: Somatosensory–ventral visual integration

The fourth cluster showed increased connectivity between somatosensory and ventral visual regions (F = 9.72, p-FDR = .046). The strongest effects were observed between the left postcentral gyrus and bilateral temporal–occipital fusiform cortex. In particular, the left postcentral gyrus showed strong coupling with the left temporal–occipital fusiform cortex (T = 5.18, p-FWE = .004) and the right temporal–occipital fusiform cortex (T = 4.89, p-FWE = .004). These were among the strongest positive gPPI effects in the analysis and indicate increased somatosensory–ventral visual coupling during sequence relative to random blocks.

## 4 Discussion

This study provides crucial information about behavioural, and physiological characteristics of MSL, as explored by the Serial Reaction Time Task (SRTT). Overall, the behavioural, BOLD activation, and connectivity findings provide a consistent picture of implicit motor sequence learning, and the suitability of the SRTT as a protocol to explore MSL in fMRI in healthy humans. Behavioural performance was stable across both sessions and task versions, with no significant effect of Session and Task Version. This indicates that the protocol is well suited for repeated-measures and crossover designs, and that the two task versions can be counterbalanced and pooled without introducing systematic bias. Three main findings should be highlighted. First, the SRTT protocol produced clear and reproducible markers of motor sequence learning. Second, sequence-specific brain activation involved a distributed network including the bilateral supplementary motor area (SMA), left superior parietal cortex, bilateral parietal operculum, ventral visual cortex, cerebellar Crus I, and the primary motor cortex (M1), although the latter not isolated as a separate sequence-specific cluster, but strongly and bilaterally engaged by the task and showing additional sequence-related activation that marks it as a central node of the execution network. Third, the seed-based connectivity (SBC) and gPPI results suggest two discernable functional components: a left-lateralised motor execution network and a mainly right-hemispheric supervisory network. Together, these findings provide an appropriate neurobiological basis for selecting targets in the planned focal-tDCS intervention.

### 4.1 Behavioural characterisation

The analyses of Session and Task Versions showed strong block-related effects across all four behavioural measures. The effect sizes for Block were large, with ηp^2^ values ranging from .14 to .55. In contrast, there were no significant effects of Session, Task Version, or their interaction. The expected slowing in the random blocks, as compared to the sequence blocks, was observed. Performance in B6 and B9 did not differ from B1 for either raw or normalised reaction time. This return to slower responses during random blocks is a typical behavioural marker of implicit sequence learning, because it separates sequence-specific learning from general practice-related improvement (Nissen **&** Bullemer, 1987; Robertson, 2007; Schwarb **&** Schumacher, 2012).

The fact that this pattern was observed across two sessions and two task versions supports the use of the SRTT as a reliable group-level outcome measure in the upcoming tDCS phase. It also suggests that Task V2 and Task V3 can be randomised, or counterbalanced without introducing a systematic bias in the block effect, and that results can be pooled for analysis.

At the same time, the test–retest reliability results are consistent with the known reliability paradox in cognitive and motor tasks (Hedge et al., 2018; Oliveira et al., 2023, 2024). Reaction times in the random blocks were highly stable across sessions, with ICC (2,1) values of approximately .80–.85. In contrast, reaction times in the sequence blocks showed only modest retest-reliability, with ICC (2,1) values of approximately .29–.53. This difference is meaningful. Random blocks mainly reflect stimulus-driven motor speed, which appears to be a relatively stable individual characteristic. Sequence blocks, however, capture an ongoing learning process, and the learning trajectory can vary between sessions.

The ICC (3,1) values were only slightly higher than the ICC (2,1) values, indicating that average session-related shifts were small. For the active tDCS phase, these findings suggest that the SRTT might mainly be used as a robust group-level measure of procedural learning rather than as a stable individual-difference measure. However, reaction time in the random blocks may still provide a useful and stable measure of general motor speed.

### 4.2 A sequence-specific multi-system activation network

For the comparison of both SEQ and RND versus resting state, the latter engaged a largely overlapping motor-execution network, including the bilateral primary motor and somatosensory cortex, the SMA, the dorsal premotor cortex (on the precentral and superior frontal gyri), the central and parietal operculum, and cerebellar lobules VI and VIII, with very similar peak intensities.

This convergence is expected, because the overt motor demand of responding to a visual cue with a button press is identical in both conditions, and indicates that a common cortico–striato–cerebellar execution network is recruited regardless of whether the cue order is predictable (Hardwick et al., 2013; Hikosaka et al., 1996; Doyon & Benali, 2005). Direct comparison of the two block types against a shared resting state therefore conflates execution-related and sequence-specific processes; the sequence-specific component is isolated only by the direct SEQ > RND contrast discussed below. A notable feature of both baseline contrasts was the strong, bilateral engagement of the primary motor cortex (M1). Although M1 is often regarded primarily as an execution structure, converging evidence indicates that it also contributes to sequence-specific learning and its consolidation, on two fronts. First, non-invasive brain-stimulation studies show that anodal tDCS over M1, but not over premotor or prefrontal control sites — facilitates implicit sequence learning, and in our own work M1 stimulation selectively improved sequence but not random stimuli performance (Nitsche et al., 2003; Reis et al., 2009; Stagg et al., 2011). Second, newer imaging studies support a critical role of M1 specifically in early sequence learning: network-based and multivariate fMRI report sequence-specific representations and a dynamic reconfiguration of M1 cortical coupling as the sequence is acquired (Hamano et al., 2020; Wiestler & Diedrichsen, 2013), and a recent meta-analysis confirms consistent M1 activation in early MSL (Chen et al., 2026), although some recent fMRI work has questioned the specificity of this contribution (Berlot et al., 2020; Jäger et al., 2022). M1 is assumed to encode primarily effector-specific information, while premotor, parietal, and supplementary motor areas represent the more abstract sequence, independent of the effectors used (Grafton et al., 1998). Consistent with a genuine learning role, the direct SEQ > RND contrast in the present data did not merely reproduce the shared execution component: left M1 (precentral gyrus) showed significantly greater activation during sequence than random blocks (peak T = 5.50), within a large left fronto-parietal cluster that also included the premotor and superior parietal cortex. M1 was, as expected, strongly active engaged in both block types, but this additional sequence-related increase indicates genuine sequence-specific involvement rather than execution alone. Taken together with the stimulation and imaging literature, this robust and partly sequence-specific M1 activation supports M1 as a mechanistically reasonable target for the planned focal-tDCS study phase (see Section 4.6).

The six clusters that survived correction in the SEQ > RND contrast closely match brain regions previously reported in activation-likelihood meta-analyses of motor sequence learning (Hardwick et al., et al., et al., 2013; Janacsek et al., 2020). Together, these clusters can be understood as parts of a distributed sequence-learning network.

The largest cluster was observed on the left superior parietal lobule. This region may support the spatial component of sequence learning by helping to maintain and update expectations about the location of the next cue (Willingham et al., 2002; Wiestler & Diedrichsen, 2013). Two additional clusters were located in bilateral medial premotor regions, including the left and right SMA/cingulate motor area. These findings are consistent with the well-established role of the SMA in implicit motor sequence learning. The SMA may support motor planning and the organisation of individual movements into larger sequence units. Its bilateral involvement also fits with the concept that the SMA represents sequential structure beyond the specific hand used for the response (Nachev et al., 2008; Shima & Tanji, 2000).

Two further clusters were found in bilateral parietal opercular regions, corresponding to the secondary somatosensory cortex. These regions may reflect predictive somatosensory processing. In other words, when the upcoming movement can be predicted, the brain may anticipate the sensory consequences of the next finger press. Finally, the left ventral visual cluster included the lingual gyrus, fusiform gyrus, lateral occipital cortex, and left cerebellar Crus I. This cluster may reflect prediction of the upcoming visual cue stream.

The involvement of cerebellar Crus I is consistent with the role of the cerebellum in cognitive aspects of early motor learning (Bernard & Seidler, 2013; King et al., 2019; Stoodley & Schmahmann, 2009). Rather than encoding the sequence itself, Crus I may contribute to predictive, error-based processing of the upcoming visual cue stream, in line with the proposed role of the cerebellum in forming internal forward connections during early learning rather than in storing the sequence representation.

### 4.3 Connectivity dynamics: a non-monotonic SBC trajectory

The SBC analysis identified a strongly connected cortico–striato–cerebellar motor network in all task conditions. Importantly, this network was already present in the initial random condition, before participants had been exposed to the sequence. This suggests that the network reflects a general motor execution system rather than a purely sequence-specific network. Motor sequence learning therefore appears to reorganise an existing motor network rather than recruit a completely new one.

Within this motor network, the L_M1–L_SMA connection was the strongest, or joint-strongest, connection across all five conditions. This is consistent with the classical premotor–motor pathway (Dum **&** Strick, 1991; Hikosaka et al., 1996; Honda et al., 1998). This connection may therefore represent a central task-general motor pathway on which sequence-related changes are built.

The cluster-level F-statistic followed a non-linear pattern across conditions, with overall network coupling falling from the initial random baseline to early sequence exposure, rising again through the middle phase, declining during late learning, and reaching its maximum during the interleaved random blocks. This pattern may reflect three overlapping learning-related processes. First, the drop from the initial random block to early sequence learning may reflect a temporary redistribution of cognitive resources. On first exposure, attention and working memory may be engaged in processing the unfamiliar cue sequence, reducing coupling within the core motor network. This is consistent with previous reports of decreasing task-related activation as learning proceeds (Berlot et al., 2020).

Second, the activation increases from early to middle sequence learning may reflect the strengthening of an emerging sequence representation. Connections that were weak or absent during early learning became stronger as participants acquired the sequence. The clearest example was the L_SMA–L_Putamen connection. This connection was not significant at the initial random Block 1, became weakly significant during early sequence learning, and then increased during later learning phases. This pattern is consistent with suggestions that cortico-striatal connections are gradually shaped during early sequence learning (Tzvi et al., 2014, 2015).

Third, the decrease from middle to late sequence learning may reflect the beginning of automatisation. At this stage, L_M1–L_Putamen coupling fell below threshold, L_M1–R_Cerebellum coupling became weaker, and the SMA became the main cortical interface with subcortical motor regions. This shift is in line with previous suggestions of motor learning and automatisation (Bassett et al., 2015; Lehéricy et al., 2005; Steele & Penhune, 2010; Berlot et al., 2020; Ölveczky, 2022).

A complementary interpretation applies to the connections that re-emerged in the late and interleaved phases. The L_SMA–R_Cerebellum coupling, which was weak or absent during early and middle sequence learning, became significant again during late sequence learning and was strongest during the interleaved random blocks, while the L_SMA–L_Putamen coupling continued to strengthen across the same phases. Because the interleaved random blocks violate the expectation built up from the learned sequence, this late re-engagement may reflect expectation- or prediction-error processing rather than further sequence consolidation. Such a violation could also exert a delayed influence that carries into the adjacent late sequence blocks, enhancing cerebellar involvement in particular, in line with the role of the cerebellum in encoding sensory prediction and expectation errors (see Section 4.2).

A particularly important SBC finding was that network coupling was stronger during the interleaved random condition (F = 72.55) than during the initial random cue condition, even though both conditions used a pseudo-random stimulus order. This suggests that the two random conditions were not functionally identical. In the initial baseline block, participants responded to random stimuli before learning any sequence structure. In the later interleaved random blocks, however, participants had already formed expectations based on the learned sequence. When the expected sequence was interrupted by random stimuli, participants likely had to suppress the learned prediction and return to feedback-driven sensorimotor control. This may explain the stronger moment-to-moment coupling within the motor network during the interleaved random condition (Shadmehr **&** Krakauer, 2008; Shin & Ivry, 2003).

### 4.4 Task-modulated connectivity: a right-hemisphere supervisory overlay

In contrast to the SBC analysis, which characterised a predefined motor network separately for each block, the gPPI analysis was exploratory and addressed the whole brain: through the SEQ > RND contrast it tested task-modulated connectivity across all atlas regions, directly isolating sequence-specific connections — including connections outside the predefined motor network - but without the block-specific temporal resolution that the SBC provided.

The gPPI analysis identified four functional clusters in which connectivity differed between sequence and random blocks. The most prominent feature was the dominance of right-hemispheric connectivity effects. Although this may appear to differ from the left-dominant BOLD activation pattern, the two findings are not contradictory. BOLD activation reflects local processing demands, which are expected to be strongest in the contralateral left hemisphere during a right-hand motor task. Functional connectivity, however, reflects coordination between brain regions. A region can therefore act as an important connectivity hub even if it is not the site of strongest local activation (Di et al., 2018; Friston et al., 1997).

The gPPI and SBC analyses are complementary because they test different questions rather than because they differ in sensitivity. SBC quantifies the average coupling of the predefined motor network within each task phase, whereas gPPI tests whether that coupling is modulated by the psychological context (SEQ vs RND) through a psychophysiological interaction term. Because the two approaches model the data differently average within-condition coupling versus a condition-by-seed interaction they highlight partly different connections and are therefore best interpreted together rather than as redundant analyses.

A further distinction is temporal in origin: SBC was estimated separately for each task phase and therefore preserved the block-wise temporal dynamics of learning across the session, whereas gPPI collapses across the whole time series and lacks this temporal resolution. Because connectivity in the SRTT evolves over the course of learning (Bazin & Steele, 2022; Steele & Penhune, 2010), this loss of temporal information may attenuate some sequence-related connections in the gPPI analysis, so that the two approaches are best read as complementary.

The right-hemispheric dominance of sequence-related connectivity is consistent with the role of the right hemisphere in visuospatial attention and spatial sequence monitoring (Corbetta & Shulman, 2002, 2011; Vossel et al., 2014). It is also consistent with evidence that right-lateralised supervisory and executive processes contribute to motor sequence learning (Grafton et al., 2002; Serrien et al., 2006).

The four gPPI clusters indicate that sequence-related connectivity is organised along dorsal and ventral processing routes. Cluster 1 involves a dorsal visuospatial–motor pathway, connecting bilateral visual areas with right sensorimotor and parietal regions. This pathway may help participants track the spatial position of the cues and translate this visual information into the appropriate motor response during sequence learning. Cluster 4 appears to involve a ventral somatosensory–visual integration pathway. This cluster showed connectivity between the left postcentral gyrus and bilateral fusiform cortex. This pattern may reflect the integration of sensory feedback from the responding hand with visual information about the cue identity and location.

Cluster 3 showed reduced coupling during sequence blocks between left SMA and left M1, and between left central opercular cortex and left M1. This pattern may reflect increasing automatisation. As the sequence becomes more familiar, performance may rely less on effortful SMA–M1 coordination and more on more efficient motor execution (Bassett et al., et al., et al., 2015; Lehéricy et al., 2005; Steele & Penhune, 2010). Finally, Cluster 2 reflected a parieto-premotor–thalamic integration network. This network overlaps with the classical cortico–striato–thalamo–cortical circuit that has been linked to procedural sequence learning (Alexander et al., 1986; Doyon & Benali, 2005; Hikosaka et al., 2002).

### 4.5 Integration: a two-component architecture for SRTT performance

Taken together, BOLD activation, SBC, and gPPI findings suggest a two-component network relevant for SRTT performance. These results should therefore be interpreted as complementary levels of the same learning system rather than as separate findings.

The first component is a left-lateralised cortico–striato–cerebellar motor execution network. This network supports button-press responses and is active across all task conditions. It is reflected in the left-dominant SEQ > RND BOLD activation pattern and in the SBC results, where a strongly connected motor network was already present in Block 1. Within this network, the L_M1–L_SMA connection appears to a central, largely task-general core pathway (Doyon & Benali, 2005; Hardwick et al., 2013; Hikosaka et al., 2002). We describe this core as largely, rather than purely, task-general. Although it is engaged across both predictable and unpredictable blocks, stimulation studies indicate that nodes within it — in particular M1 and SMA — also contribute to sequence-specific learning. Stimulation studies demonstrate a causal role (Nitsche et al., 2003; Reis et al., 2009; Stagg et al., 2011) and also imaging studies report sequence-specific representations in these regions (Grafton et al., 1998; Honda et al., 1998; Wiestler & Diedrichsen, 2013; Hamano et al., 2020; Chen et al., 2026).

The second component is a mainly right-hemispheric supervisory network that becomes more strongly engaged when the cue order is predictable. This network links visual regions with right sensorimotor, parietal, SMA, and thalamic areas. Its right-sided dominance is consistent with the role of the right hemisphere in visuospatial attention and the monitoring of spatial sequences (Corbetta & Shulman, 2002, 2011; Grafton et al., 2002; Serrien et al., 2006).

The two connectivity analyses also show converging learning-related changes. The reduced left SMA–M1 coupling observed in the gPPI analysis, together with the middle-to-late reorganisation seen in the SBC analysis, suggests a gradual shift from effortful, M1-centred control toward more internally guided, SMA-based motor control as learning progresses (Bassett et al., 2015; Lehéricy et al., 2005; Steele & Penhune, 2010; Berlot et al., 2020; Ölveczky, 2022).

At the same time, the visual and somatosensory BOLD activations, together with the gPPI findings, suggest increasing perceptual prediction and haptic–visual integration. Visual prediction is reflected in the left ventral visual activation and in the visual-to-sensorimotor coupling observed in gPPI Cluster 1. Haptic–visual integration is reflected in the bilateral S2 activation and in the postcentral-to-fusiform coupling observed in gPPI Cluster 4 (Lacey **&** Sathian, 2014).

Overall, the three analyses provide complementary information. BOLD activation identifies which brain regions are engaged during sequence learning. SBC shows how the core motor network changes across different learning phases. gPPI shows how connectivity differs between sequence and random blocks.

### 4.6 Dissociation between group-level effects and individual-level reliability

A consistent topic across the behavioural, activation, and connectivity analyses is a dissociation between group-level effects and individual-level reliability. At the group level, the SRTT produced robust, reproducible effects that were stable across the two sessions and the two counterbalanced task versions; paired t-tests and ANOVAs revealed no systematic session or version differences. At the individual level, by contrast, test–retest reliability was only low to moderate, and for the SEQ > RND difference it approached zero.

This dissociation is the expected consequence of the well-known reliability paradox (Hedge et al., 2018). The intraclass correlation expresses between-subject variance relative to total variance, so when a paradigm evokes a near-universal effect, as the SRTT does for sequence learning, between-subject variance and consequently the ICC is small even though the measurement is valid and the group-level effect is stable. Difference measures such as SEQ > RND compound this, because subtracting two highly similar conditions cancels their shared signal and roughly doubles the relative noise. Low reliability of this kind is therefore a property of the paradigm and the contrast, not evidence of unstable measurement.

For this reason, our decision to pool the two sessions and the two task versions was based on the group-level equivalence established by the t-tests and ANOVAs, and not on the ICC values. Reporting both levels makes the logic explicit: the group-level analyses allow pooling and support the use of the SRTT as a robust outcome measure for the planned focal-tDCS phase, whereas the reliability analyses delineate the limits of individual-level inference. Accordingly, in the stimulation phase the SRTT and its imaging correlates are best interpreted as group-level markers of motor-sequence learning, with individual-level change treated cautiously.

### 4.7 Implications for the focal tDCS phase

The two-component network identified in this study has direct implications for the design of the upcoming active-tDCS phase. Anodal tDCS over the left M1, a standard target for motor-learning interventions, is expected to act primarily on the left-lateralised motor execution network identified by the BOLD and SBC analyses.

Its effects should therefore be detectable as changes in within-network SBC coupling, especially in the balance between L_M1 and its cortical and striatal connections with L_SMA and L_Putamen, which changed systematically across the different phases of sequence learning in the present study. Beyond its practical accessibility, M1 is a content-driven choice on three grounds. First, it was among the most strongly and reliably activated regions in both the SEQ > Baseline and RND > Baseline contrasts, confirming that it is robustly engaged by the task in the present protocol. Second, M1 is a hub of the left-lateralised execution network and a node of the L_M1–L_SMA pathway that showed the most consistent coupling across all learning phases, so stimulation here is well placed to modulate the network dynamics we observed. Third, M1 has an established causal role in motor-sequence acquisition and consolidation in tDCS and TMS studies (Muellbacher et al., 2002; Nitsche et al., 2003; Reis et al., 2009; Stagg et al., 2011), making it the most evidence-based single target for focal NIBS intervention. In our own 2003 study, stimulation of control sites (premotor and prefrontal cortex) during early learning was without effect, underlining the specificity of M1, and its contribution is further supported by a range of imaging studies (Honda et al., 1998; Hamano et al., 2020; Chen et al., 2026). M1 is moreover suited in particular to modulating the early-learning stage, whereas premotor regions are more relevant to later reconsolidation (Nitsche et al., 2003, 2010). Together, these grounds make M1 the most evidence-based single target for focal intervention.

Two methodological points are also important for the upcoming phase. First, group-level effects should be treated as the main behavioural outcome, because individual-difference measures of sequence-block performance showed poor to modest reliability (Hedge et al., 2018; Oliveira et al., 2023). Second, the SBC analysis tracks how each connection changes across the different learning stages, from early to late. This gives a step-by-step picture of the network within a single session, so a stimulation effect can be linked to a specific connection at a specific stage of learning. A single activation contrast alone is not well suited for this, because it is based on an average over the whole session.

## 5 Limitations

Several limitations should be considered when interpreting these findings. First, the sample size was modest by current fMRI standards (N = 20). As a result, more subtle connectivity and activation changes associated with the middle-to-late reorganisation of the network did not reach significance and should be confirmed in larger samples.

A further limitation is the reliance on fMRI alone. We chose fMRI for its high spatial resolution, which is well suited to localise task-specific activations while still capturing the time course of learning across blocks. Complementary methods such as MEG or PET could nonetheless add relevant information that fMRI cannot. MEG would resolve the fast, trial-by-trial dynamics of the self-paced SRTT, while the temporal resolution of the BOLD signal and gPPI capture these temporal dynamics only partially. Furthermore, MEG would allow to identify oscillatory markers of motor learning, such as beta-band activity, to be tracked. PET, with an appropriate radiotracer, is suited to probe neurochemical processes relevant to learning that fMRI cannot assess. Combining these methods with the present fMRI findings would therefore validate and extend them rather than simply replicate these. Finally, because the SRTT was self-paced, the blocks followed one another without fixed, timed rest intervals; adding a 20–30 s rest between blocks in future studies would improve the separation of block-related responses in the GLM and strengthen the activation and connectivity estimates.

## 6 Conclusion

The present SRTT protocol produced robust and reproducible group-level markers of implicit motor sequence learning across two sessions and two independent task versions. At the same time, the behavioural data reproduced the well-known dissociation between group-level and individual-level reliability.

Beyond the specific neural findings, the present study illustrates a methodological point that will be important for the stimulation phase: across behaviour, BOLD activation, and connectivity, the SRTT showed a consistent dissociation between stable, reproducible group-level effects and low-to-moderate individual-level reliability, the latter being an expected consequence of the reliability paradox rather than a sign of unstable measurement. Because the two sessions and the two task versions did not differ at the group level, the data could be validly pooled on that basis, and the SRTT is best used as a robust group-level marker of procedural learning rather than as an individual-difference measure.

For physiological correlates of SRTT performance, fMRI revealed sequence-specific activation distributed across a broad learning-related network, including the bilateral SMA and dorsal premotor cortex, left superior parietal cortex, bilateral parietal operculum, ventral visual cortex, and cerebellar Crus I. The primary motor cortex, although not isolated by the sequence-specific analysis, was strongly and bilaterally engaged by the task itself and remains together with the SMA and premotor cortex a key node of the execution network and the primary stimulation target for the next phase. The connectivity analyses further revealed a coherent two-component functional architecture: a left-lateralised cortico–striato–cerebellar motor-execution network, whose configuration shifted non-linearly across the session in a manner consistent with successive phases of resource redistribution, consolidation, and automatisation; and a predominantly right-hemispheric visuospatial network that was more strongly engaged as the stimulus sequence became predictable.

Together, these findings establish an empirical foundation for interpreting the upcoming active focal-tDCS phase of the programme, and identify specific within-network connectivity markers that can be used to test whether M1-tDCS shifts the motor network toward a more consolidated and automatised pattern of learning.

## Acknowledgments

This research was funded by the German Research Foundation (DFG) (Research Unit 5429/1 (467143400), NI 683/17-1).

## Ethics Statement

The study was conducted in accordance with the Declaration of Helsinki and approved by the IfADo Ethics Committee under approval number 217, dated 10 May 2022. All participants gave written informed consent before participation and were free to withdraw at any time without consequence.

## Data Availability Statement

This study was pre-registered on the Open Science Framework (OSF). The pre-registration protocol is available under OSF Registries (https://osf.io/t37u2). The data that support the findings of this study are available upon request from the corresponding author. The data are not publicly available due to ethical restrictions related to the study participants.

