## Supplementary Material for "Determining Reliable Neural Targets for enhancing Motor Sequence Learning: A preparatory study for focal tDCS"

**Group-level reproducibility (t-tests) and individual-level reliability (ICC) of SRTT activation and connectivity**

Because the SRTT elicits a robust learning effect, individual differences are small — the condition under which the reliability paradox (Hedge et al., 2018) predicts low test–retest reliability despite an entirely valid measurement. We therefore characterise reliability at two levels: group-level reproducibility of the effect (paired t-tests / repeated-measures ANOVA) and individual-level stability (ICC). The dissociation between strong group effects and poor-to-moderate ICCs indicates that low ICCs reflect the statistical nature of the paradigm rather than genuine session-to-session differences, and that the SRTT is appropriately evaluated at the group level — a pattern observed consistently across behaviour and fMRI.

***Session labels.*** *“Session 1” and “Session 2” denote the two task-fMRI sham sessions (≈ 1 week apart). Group comparisons are within-subject (paired).*

### **S1. Activation**

#### **S1.1 Session reproducibility (paired t-tests, Session 1 vs 2)**

Whole-brain SPM paired t-tests comparing Session 1 with Session 2 within subject (p < .001 uncorrected, cluster-level FWE correction at p < .05) were conducted. No cluster survived correction for any contrast; the largest sub-threshold cluster and its statistics are reported to show how far the effects are from significance.

| **Contrast (Session 1 vs 2)** | **Largest cluster (k)** | **cluster-pFWE** | **peak T** | **Outcome** |
| --- | --- | --- | --- | --- |
| **SEQ vs baseline** | 52 vox | 0.51 | 4.32 | No session difference |
| **RND vs baseline** | 15 vox | 0.96 | 3.88 | No session difference |
| **SEQ > RND** | ≤ 35 vox | ≥ 0.76 | ≤ 5.14 | No session difference |

In the Session 1 > Session 2 direction no clusters were present for any contrast (for SEQ > RND, none even at the uncorrected threshold); the values above represent the largest cluster in the Session 2 > Session 1 direction. No cluster approached significance (all cluster-pFWE ≥ .51, all peak-pFWE ≥ .49), indicating that task activation does not differ between sessions and that Session 1 and Session 2 can be pooled.

Positive equivalence evidence (a-priori ROIs). Because a non-significant difference is not proof of equivalence, the Session 1 vs 2 comparison was repeated at 11 a-priori ROIs with a JZS Bayes factor (BF₀₁ > 3 = moderate evidence for no session difference; < 1 = evidence for a difference). No ROI differed between sessions after correction (paired t).

| **ROI** | **SEQ BF₀₁** | **RND BF₀₁** |
| --- | --- | --- |
| L_M1 | 3.10 | 4.21 |
| SMA | 3.62 | 4.20 |
| L_PMd | 1.98 | 2.88 |
| L_S1 | 2.89 | 4.15 |
| L_Parietal | 2.23 | 3.12 |
| L_Putamen | 4.01 | 4.18 |
| L_Caudate | 3.66 | 4.05 |
| L_Thalamus | *0.34* | *0.80* |
| R_Cerebellum V | *0.94* | 1.23 |
| R_Cerebellum VI | *0.66* | *0.73* |
| R_Cerebellum VIII | 2.48 | 3.01 |

Most ROIs favour no session difference (BF₀₁ ≈ 2–4), supporting pooling. The exceptions — thalamus and cerebellar lobules V/VI (BF₀₁ < 1, italicised) — show weak evidence of a small session shift; in the paired t-tests only L_Thalamus (SEQ p = .020) and R_Cerebellum VI (SEQ p = .046) were nominally significant, but neither survived correction across the 11 ROIs. These effects are thus minor and do not affect the whole-brain conclusion.

#### **S1.2 Task-version comparison (paired t-tests, V2 vs V3)**

The same whole-brain paired t-tests comparing Task V2 with Task V3 (within subject; p < .001 uncorrected, cluster-FWE p < .05). Unlike sessions, versions were not fully equivalent: one SEQ cluster survives cluster-level correction.

| **Contrast (V2 vs V3)** | **Largest cluster (k)** | **cluster-pFWE** | **peak T** | **Outcome** |
| --- | --- | --- | --- | --- |
| **SEQ vs baseline** | 172 vox @ −60,14,20 | 0.019 | 5.53 | One cluster differs (L IFG / ventral premotor) |
| **RND vs baseline** | 124 vox | 0.092 | 5.43 | No version difference (n.s.) |

Only the SEQ V2 > V3 contrast yielded a surviving cluster, in the left inferior frontal / ventral premotor cortex (cluster-pFWE = .019; peak-pFWE = .46, i.e. not significant at the peak); all other version clusters were non-significant (cluster-pFWE ≥ .09). At the ROI level the version equivalence evidence is correspondingly weaker than for sessions (median BF₀₁ ≈ 1.1 for both SEQ and RND; several ROIs with BF₀₁ < 1, including left parietal and cerebellar V/VI). The two versions therefore differ modestly in left fronto-premotor cortex activation. Because Task V2 and V3 were counterbalanced across sessions, this version effect is however balanced by the session and group analyses by design.

#### **S1.3 Individual-level reliability (ICC) SEQ, RND, SEQ > RND**

Test–retest ICC shown across the two sessions (PyReliMRI), at three spatial scales (whole-image, whole-brain voxelwise, and atlas-ROI ICC). ICC(2,1) absolute agreement; ICC(3,1) consistency. ICC was near-identical for the condition maps (median gap ≈ −0.01), indicating negligible systematic between-session shift. ICC values were interpreted according to the ICC ranges poor < .50, moderate .50–.75, good .75–.90 (Koo & Li, 2016).

| **Reliability metric** | **SEQ vs BL** | **RND vs BL** | **SEQ > RND** |
| --- | --- | --- | --- |
| Whole-image I2C2 | 0.47 | 0.46 | 0.08 |
| Whole-brain voxelwise ICC (median) | 0.43 | 0.38 | 0.04 |
| Voxels with ICC > 0.40 | 54% | 46% | 4% |
| Atlas ROI ICC, 132 regions (median) | 0.53 | 0.46 | low (≈ contrast) |

**Pattern.** Both condition maps (SEQ, RND) are moderately reliable; the SEQ>RND difference map collapses on every metric (I2C2 = 0.08). Low contrast reliability is the expected difference-score result, not a data problem. Behaviourally (main text), random-block RT was the more reliable measure (ICC ≈ .80–.85) compared to sequence RT modest (≈ .29–.53); the neural and behavioural reliability orders do not necessarily coincide because they index different constructs (between-subject RT variance vs spatial map reproducibility).

### **S2. Connectivity**

#### **S2.1 Session reproducibility (paired t-tests, Session 1 vs 2)**

SBC (5-seed motor network; 10 edges) session reproducibility was tested with per-edge paired t-tests (Session 1 vs 2; FDR across the 10 edges) with TOST and JZS BF₀₁ equivalence. For gPPI (132-region atlas; SEQ>RND; 17,292 edges) per-edge t-tests are not appropriate; its session reproducibility is reported as whole-pattern I2C2 in S2.3.

SBC session reproducibility (paired t per edge, Session 1 vs 2):

| **Phase** | **Edges differing (FDR<.05)** | **Median BF₀₁** | **Outcome** |
| --- | --- | --- | --- |
| Block 1 | 1 / 10 | 3.23 | Difference Cerebellum–Putamen |
| Blocks 2&3 | 0 / 10 | 3.62 | No session difference |
| Blocks 4&5 | 0 / 10 | 3.50 | No session difference |
| Blocks 7&8 | 0 / 10 | 3.31 | No session difference |
| Blocks 6&9 | 1 / 10 | 3.69 | Difference Putamen–PMC |

Across all phases only 2 of 50 edges differed between sessions after FDR — R_Cerebellum–L_Putamen (initial RND block, p-FDR = .014) and L_Putamen–L_PMC (interleaved RND, p-FDR = .021); both involve the putamen and both fall in random-block phases. For all remaining edges BF₀₁ favoured no session difference (per-phase medians 3.2–3.7). Session reproducibility therefore strongly supports pooling the two sessions for the connectivity analyses. TOST flagged no edge as formally equivalent, consistent with it being underpowered at this sample size; BF₀₁ is the more informative equivalence statistic here.

#### **S2.2 Task-version reproducibility (V2 vs V3)**

Connectivity reproducibility across the two task versions, paired within subject using the V2/V3 counterbalanced map is shown. SBC: per-edge paired t (V2 vs V3) per phase with FDR + BF₀₁. gPPI: whole-pattern I2C2 + spatial correlation between the V2 and V3 SEQ>RND matrices.

SBC — version paired t per edge (FDR across 10 edges per phase):

| **Phase** | **Edges differing (FDR<.05)** | **Median BF₀₁** | **Outcome** |
| --- | --- | --- | --- |
| Block 1 | 0 / 10 | 3.46 | No version difference |
| Blocks 2&3 | 0 / 10 | 2.07 | No version difference |
| Blocks 4&5 | 0 / 10 | 1.77 | No version difference |
| Blocks 7&8 | 0 / 10 | 3.58 | No version difference |
| Blocks 6&9 | 0 / 10 | 3.74 | No version difference |

gPPI — version whole-pattern reproducibility:

| **Metric** | **Value** |
| --- | --- |
| Whole-matrix I2C2 (V2 vs V3) | 0.006 (null −0.003 ± 0.010; p = 0.199) |
| Group spatial correlation (r; ρ) | 0.25; 0.18 |
| Edge ICC(3,1): median; % ≥ 0.50 | 0.014; 1.8% |

No SBC edge differed between task versions V2 and V3 in any phase after FDR; BF₀₁ favoured no version difference in most edges (per-phase medians 1.8–3.7). Two edges were nominally significant before correction (Putamen–M1 in early sequence, Putamen–PMC in late sequence) but did not survive FDR. For gPPI, the SEQ>RND difference pattern shows the same near-zero individual-level reliability as for sessions (I2C2 indistinguishable from null), while the group-average pattern is moderately consistent across versions (spatial r = 0.25). Connectivity therefore does not show the left fronto-premotor version effect seen in activation (S1.2): at the motor-network level the two versions are reproducible, so pooling the counterbalanced versions is well supported for the connectivity analyses.

#### **S2.3 Individual-level reliability (ICC) SBC and gPPI**

Edgewise test–retest ICC across sessions (ICC(3,1) primary for connectivity, ICC(2,1)

SBCedgewise ICC by phase (ses 1 vs 2):

| **Phase** | **Blocks** | **Median ICC(3,1)** | **Median ICC(2,1)** | **Edges ≥ 0.50 (ICC3)** |
| --- | --- | --- | --- | --- |
| Initial RND | 1 | 0.17 | 0.16 | 1/10 |
| Early SEQ | 2,3 | 0.00 | 0.00 | 0/10 |
| Middle SEQ | 4,5 | 0.13 | 0.13 | 2/10 |
| Late SEQ | 7,8 | 0.14 | 0.15 | 1/10 |
| Interleaved RND | 6,9 | 0.07 | 0.07 | 0/10 |
| **SEQ (all)** | 2–5,7,8 | 0.26 | 0.26 | 2/10 |
| **RND (all)** | 1,6,9 | 0.22 | 0.17 | 1/10 |
| **All blocks** | 1–9 | 0.36 | 0.37 | 3/10 |

Edge-level SBC reliability is low overall (median ICC mostly 0.0–0.4), consistent with the connectivity benchmark of Noble et al. (2019), mean edge ICC = 0.29 (95% CI 0.23–0.36). Reliability increases with the number of pooled blocks: SEQ (6 blocks, 0.26) exceeds RND (3 blocks, 0.22), and the all-blocks pool (9 blocks) is highest (0.36) in accordance with the expected averaging effect (more blocks → lower within-subject noise, resulting in higher ICC). ICC(3,1) and ICC(2,1) are near-identical (median gap ≈ 0.00), indicating no systematic between-session shift. As at the activation level, the difference-style phases (early sequence) are least reliable.

gPPI — SEQ>RND whole-pattern and edgewise reliability (sessions; real values):

| **Metric** | **Value** |
| --- | --- |
| Whole-matrix I2C2 | 0.006 |
| Permutation null (mean ± SD); p | 0.000 ± 0.010; p = 0.275 |
| Edge ICC(3,1): median [IQR] | 0.008 [−0.16, 0.17] |
| Edges with ICC(3,1) ≥ 0.40 | 5.2% |
| Edges with ICC(3,1) ≥ 0.50 | 1.7% |
| Directed edges; N | 17,292; 19 |

The gPPI SEQ>RND difference shows no individual-level reliability (I2C2 indistinguishable from its permutation null). This is the expected difference-score / reliability-paradox result and is reported for completeness; the gPPI group effect (reported in the main text) remains robust.
